# Genetic differentiation and adaptive evolution of buff-tailed bumblebees in Asia

**DOI:** 10.1101/2024.08.13.607144

**Authors:** Long Su, Lele Ding, Paul H. Williams, Yan Liu, Ruijuan Wang, Xiaoyan Dai, Shan Zhao, Haolin Fu, Xiaomeng Zhao, Quangui Wang, Yancan Li, Huiling Sang, Robert M. Waterhouse, Yifan Zhai, Cheng Sun

## Abstract

Bumblebees are ecologically and economically important pollinating insects, so their declines resulting from environmental changes have received intensive attention. Understanding how environmental factors shape the genetic differentiation of natural populations and identifying the genetic basis of local adaptation will provide insights into how species may cope with environmental changes. The buff-tailed bumblebee (*Bombus terrestris*) has a wide natural distribution range and has been successfully domesticated to produce commercial colonies for greenhouse pollination. Previous population genetics studies on *B. terrestris* have mainly focused on populations in Europe; however, populations in Asia, representing the eastern side of its natural distribution, have been less thoroughly sampled. To fill this gap, we collected wild *B. terrestris* samples from Asia, as well as wild *B. terrestris* from Europe and samples from domesticated colonies. We conducted whole-genome resequencing for 77 collected *B. terrestris* workers and performed population genomics analysis. Our results revealed significant genetic differentiation (*F*_ST_= 0.076) between buff-tailed bumblebees in Europe and Asia, along with notable morphological and physiological differences. Consequently, *B. terrestris* in Asia represents a distinct new genetic resource. Demographic analysis suggested that the population size of buff-tailed bumblebees had increased during historic cold periods, confirming their cold-adapted characteristics. Selective sweep analysis identified 331 genes under selection in the genomes of Asian *B. terrestris*, likely involved in their adaptation to high ultraviolet radiation, low temperature, and low precipitation in their habitats. Our research provides insights into the population genetic structure and genetic basis of local adaptation in the buff-tailed bumblebee, which will be useful for its conservation and management.

## Introduction

Bumblebees (genus *Bombus*) are ecologically and economically important pollinating insects, with ∼250 extant species worldwide (Williams, 1998). They have long been known as excellent pollinators of greenhouse tomato crops, decreasing the cost of labor and improving the yield and quality of the fruits (Velthuis & Doorn, 2006). Today their pollination services have been extended to various other crops grown in poly-tunnels and open fields (Velthuis and van Doorn 2006; Garibaldi et al., 2013; Martin et al., 2019). Bumblebees are also ecologically important pollinators, serving as the sole or predominant pollinators of large numbers of wild plants (Fontaine et al. 2006; Goulson et al. 2008).

However, the declines of bumblebees in abundance and distribution, caused by global and local environmental changes, pose a threat to both wild plants and crop pollination (Williams et al., 2009; Cameron et al., 2020). This decline has raised concerns regarding the implications for bumblebees, food security, and ecosystem stability (Bartomeus et al., 2013; Cameron and Sadd 2020; Soroye et al. 2020). Understanding how environmental factors shape the genetic differentiation of natural populations and identifying the genetic basis of local adaptation will provide insights into how species respond to environmental changes, which is fundamental to bumblebee conservation and breeding (Hoffmann and Sgrò, 2011; Franks and Hoffmann, 2012; Savolainen et al., 2013; Sang et al., 2024).

The buff-tailed bumblebee, *Bombus terrestris* (Linnaeus, 1758), is one of the most widely used pollinator species for greenhouse crop pollination, and has been artificially reared by commercial companies since the 1980s (Velthuis and Van Doorn 2006; Goulson, 2010). *Bombus terrestris* is abundant in the West Palearctic regions, with a natural distribution that includes Atlantic islands, North Africa, Mediterranean islands, continental Europe, the south Ural Mountains, and extending east to the Altai Mountains (as far east as Mongolia) (Estoup et al., 1996; Rasmont et al., 2008; Williams et al., 2012). Therefore, *B. terrestris* is a broadly distributed species and exposed to diverse climatic conditions, which makes it a good model to study population differentiation and local adaptation.

Several studies have been performed to understand the population genetic structure and differentiation of *B. terrestris*, mainly focused on European and nearby Mediterranean and Atlantic Island populations (Moreira et al., 2015; Kraus et al., 2009; Lecocq et al., 2013; Estoup et al., 1996; Widmer et al. 1998; Silva et al., 2020; Colgan et al., 2022). However, *B. terrestris* populations in Asia, representing the eastern edge of its global natural distribution, have been sampled only for morphological and COI barcode variation analyses (Williams et al., 2012).

To address in this gap, we collected wild *B. terrestris* samples from 11 Asian geographic populations, located an average distance of 5,100 kilometers from mainland Europe. Wild samples from Europe and domesticated samples from a commercial company were also collected. Whole-genome resequencing was performed on these samples, generating data for population genomic analyses to infer the population genetic structure and demographic history of *B. terrestris* and to identify candidate genes involved in the local adaptation to its Asian habitats.

## Materials and Methods

### Sample collection

Buff-tailed bumblebee (*B. terrestris*) samples were collected from both Asia and Europe for whole-genome resequencing. More specifically, 41 wild samples were collected from 9 geographic sites in Xinjiang, China, with an average distance of 254 kilometers between each site (Table S1). Additionally, 12 wild specimens were obtained from the Natural History Museum, London, United Kingdom, which originated from 6 European countries (France, Germany, Sweden, Switzerland, Turkey, England) and 3 Asian regions (Tajikistan, Kyrgyzstan, Russia (82.93N 55.01E)) (Table S1). Moreover, 24 commercial (domesticated) *B. terrestris* samples were also used. These samples were purchased from KOPPERT, a Netherlands-based company that has been involved in the breeding and domestication of buff-tailed bumblebees for over 35 years.

### Phenotypic characterization

Phenotypic traits of *B. terrestris* were measured using a stereomicroscope and the Micro Measurement and Data Analysis System (v1.01). These traits included proboscis length, forewing length (FL), forewing width (FB) , cubital vein A, cubital vein B, wing vein angles (A4, B4, D7, E9, G18, J10, J16, K19, L13, N23, O26, new angle), hooks on the right hind wing, femur length (Fe) of the right hind leg, tibia length (Ti) of the right hind leg, metabasitarsus length (ML), metabasitarsus width (MT), length of the third abdominal sternum (S3), width of the third abdominal sternum, width of the fourth abdominal sternum, length of the genitalia, and width of the genitalia.

To measure hair density on the face and mesothorax of buff-tailed bumblebees, we selected three representative areas of approximately 0.4 mm² and counted the puncture density in each area. In some cases, particularly for specimens with high hair density or hairs that formed clumps due to handling during capture and/or preservation, hairs at their insertion points (usually indicated by a micropore on the cuticle) were counted. An additional advantage of counting micropores is that it could be applied to specimens that have lost hairs (e.g., due to aging; Bosch & Vicens, 2006; Southwick, 1985; or poor handling) to obtain a measure of the original hair coverage.

Using the SUN-V Intelligent Insect Supercooling Point Tester (SUN Company, Jinan, China), supercooling points of *B. terrestris* individuals from different sampling sites were determined. Phenotypic data were correlated with supercooling point data for association and difference analysis.

### DNA extraction

Prior to DNA extraction, all buff-tailed bumblebee samples underwent surface cleaning with alcohol. DNA extraction was performed using the QIAGEN Gentra Puregene Tissue Kit (QIAGEN, Germany) following standard protocols. Given that museum specimens may have suffered DNA damage (Dabney et al., 2013), DNA repair was conducted on DNA extracted from those samples using the NEBNext FFPE DNA Repair Mix kit (New England Biolabs, NEB). In total, 77 genomic DNA samples of buff-tailed bumblebees were successfully obtained for downstream sequencing and analysis.

### DNA barcoding analysis

To verify the identity of the samples as *B. terrestris*, a barcoding PCR experiment was conducted following the protocol outlined by Hebert et al. (2004). The PCR primers were LEPr (TAAACTTCTGGATGTCCAAAAAATCA) and LEPf (ATTCAACCAATCATAAAGATATTGG). The reaction mixture (15 μL) consisted of 10× Buffer with 15 mmol/L MgCl_2_ (1.5 μL), 10 mmol/L dNTPs (0.3 μL), each primer at 15 μmol/L concentration (1.0 μL each), Taq polymerase (5 U/μL) (0.2 μL), 25 ng/μL DNA template (2 μL), and ddH_2_O (10.725 μL).

PCR cycling conditions were as follows: initial denaturation at 95 °C for 5 minutes, followed by 33 cycles of denaturation at 94 °C for 30 seconds, annealing at 56 °C for 30 seconds, extension at 72 °C for 30 seconds, and a final extension at 72 °C for 5 minutes. The PCR products then underwent Sanger sequencing to obtain barcoding sequences. Finally, sequences of the 77 samples were submitted to the BOLD Systems website (http://www.boldsystems.org/index.php) for species identification.

### Genome resequencing and read mapping

Genomic DNA quality and integrity were assessed using a nucleic acid analyzer and agarose gel electrophoresis. DNA samples of high quality, with good integrity and minimal degradation, were randomly fragmented. Libraries for paired-end sequencing were constructed according to Illumina sequencing platform requirements. Sequencing was performed using the Illumina HiSeq 2000 sequencer, generating paired-end sequencing reads with a length of 150 base pairs (bp). Additionally, because some wild samples were museum specimens, to mitigate the effects of depurination reactions that may have occurred during storage of museum specimens, we trimmed two bases from both ends of each read using FASTQ software.

### Variant calling

Genomic sequencing data from *B. terrestris* samples were aligned to the reference genome (assembly Bter_1.0 GCA_000214255.1 (Sadd et al., 2015)) using BWA-MEM software (Li and Durbin, 2009) with default parameters. The resulting alignments were converted into BAM format using SAMtools (Li et al., 2009) and sorted accordingly. Duplicate reads were removed using the GATK MarkDuplicates module (McKenna et al., 2010). Sequencing depth and coverage were then evaluated using SAMtools and BEDtools (Quinlan and Hall, 2010). For variant calling (including Single Nucleotide Polymorphisms: SNPs and Insertions-Deletions: INDELs), the GATK tools HaplotypeCaller, CombineGVCFs, GenotypeGVCFs, and SelectVariants were utilized. The obtained variants then underwent rigorous filtering using the GATK VariantFiltration module.SNP; variant annotations were conducted using ANNOVAR (Wang et al., 2015) and SnpEff (Cingolani et al., 2012).

### Kinship analysis

High genomic homogeneity among individuals from the same colony or closely related bees can lead to errors in population genomics analysis. To address this, we employed the KING-robust method (Manichaikul et al., 2010) using SNP information across the whole genome to compute pairwise kinship coefficients among the 77 samples. Since in most bumblebees, queens mate once with a single male while males can mate with multiple queens (Boomsma and Ratnieks, 1996; Nasei et al., 1998; Schmid-Hempel and Schmid-Hempel, 2000), we applied such parameters in the KING software. Individuals with kinship coefficients greater than 0.354 were considered full siblings, while those with coefficients between 0.177 and 0.354 were considered half-siblings. Therefore, during sample filtering, individuals with kinship coefficients greater than or equal to 0.177 were treated as belonging to the same colony, and only one individual per colony was retained for subsequent analyses. This resulted in a total of 67 individuals being retained for downstream analysis.

### Population genetic structure analysis

To construct the phylogenetic tree of *B. terrestris* samples, we used *Bombus ignitus*, which belongs to the same subgenus as *B. terrestris*, as an outgroup (Williams et al., 2008). The *B. ignitus* data were downloaded from the National Center for Biotechnology Information (NCBI) (PRJNA659133, (Sun et al., 2020)). Genome-wide SNP data from 67 *B. terrestris* samples and one *B. ignitus* sample were turned to phylip files using Plink software (Chang et al., 2015). The data was then used to build a Maximum Likelihood tree with RAxML (Randomized Axelerated Maximum Likelihood) software (Stamatakis, 2014). Bootstrap analysis with 1,000 replicates was performed, and a RAxML_bestTree was generated. The final phylogenetic tree was visualized using FigTree software (http://tree.bio.ed.ac.uk/software/figtree/).

Principal component analysis (PCA) was conducted on genome-wide SNP data from 67 *B. terrestris* samples. The data were first formatted into ped and map files using Plink software. PCA analysis was performed using the smartPCA module within EIGENSOFT software (Patterson et al., 2006). The results were visualized using scatter plots generated with the ggplot2 package in R.

Using structure analysis, we employed a maximum likelihood estimation algorithm assuming Hardy-Weinberg equilibrium. Structure assigns *B. terrestris* samples to distinct populations or admixed groups based on their genetic variation. This approach offers insights into the genetic composition and admixture patterns across various values of K, shedding light on the evolutionary history of *B. terrestris* populations. The results from the Structure analysis were visualized using the ggplot2 package in R, aiding in the interpretation of clustering patterns and genetic admixture among individuals.

### Gene flow tests

The ABBA-BABA test, also known as a D-test (D-statistics), is a widely used method in population genetics to detect significant deviations caused by admixture events. To examine gene flow between Asian wild populations, European wild populations and the commercial (domestic) populations of *B. terrestris*, the ABBA-BABA test was employed, with the use of *B. ignitus* as the outgroup. The analysis included two main combination models: ((Wild-Europe, Domestic), Wild-Asia, Outgroup) and ((Domestic, Wild-Europe), Wild-Asia, Outgroup). These models were cross-validated to ensure the robustness of the results. Z-scores with absolute values greater than 3 indicate statistically significant gene flow.

TreeMix analysis using whole-genome allele frequency data was then used to infer population splits and admixture events (Pickrell and Pritchard, 2012). To detect gene flow among different populations of *B. terrestris*, TreeMix analysis was also employed on the 67 *B. terrestris* samples, using *B. ignitus* as the outgroup. Wild populations of *B. terrestris* were grouped by country of collection (France, Germany, Sweden, Switzerland, Turkey, United Kingdom, Tajikistan, Kyrgyzstan, Russia, and China) to examine gene flow patterns between countries.

### Pairwise sequentially Markovian coalescent (PSMC) analysis

The PSMC model (Li and Durbin, 2011) is a method used to estimate the time to the most recent common ancestor by assessing the density of heterozygous sites across a diploid genome. In our study, we utilized the PSMC model to estimate historical population changes in Asian and European populations. Given the stringent requirements of this analysis for sequencing depth (>20×) and extensive coverage, low-quality sequencing data may introduce inaccuracies. To ensure robust outcomes, we utilized 41 high-quality resequencing datasets for *B. terrestris* samples collected from Xinjiang of China to represent the Asian population and 14 high-quality resequencing datasets from commercial (domesticated) *B. terrestris* samples bought from one European company to represent the European population.

To correlate the historical dynamics of *B. terrestris* populations with past climatic conditions, we analyzed how the population size of bumblebees was influenced by climatic changes. We employed mass accumulation rates (MAR) of loess deposits from the Loess Plateau over the past three million years as a proxy for global historical temperature changes (Sun and An, 2005). MAR values serve as indicators of global climate patterns: while lower MAR values correspond to warm and humid climates, higher values indicate cold and arid conditions.

### Analysis of climatic data

We analyzed climatic data from sampling points of 15 wild Asian bumblebees and 9 wild European bumblebees based on their geographic coordinates. Using ArcMap (version 10.2), climatic data for each sampling point were obtained from the WorldClim database (https://www.worldclim.org/). The main climatic variables included minimum temperature, annual mean temperature, maximum temperature, precipitation, elevation, and solar radiation. All climatic data were categorized into two groups based on geographic coordinates: Asia and Europe. Subsequently, we used R to conduct Wilcoxon rank-sum tests on each climatic variable to assess if there were any significant differences between the Asian and European population locations.

### Detection of positive selection in Asian *B. terrestris* populations

41 *B. terrestris* specimens collected from Xinjiang, China and three specimens collected from Tajikistan, Kyrgyzstan, and Russia represent the Asian population (collection sites in red color in Figure 1A). The European population included 9 *B. terrestris* specimens collected from France, Germany, Sweden, Switzerland, Turkey, and the United Kingdom (collection sites in green color in Figure 1A), along with one individual of domesticated *B. terrestris* bought from a European company.

**Fig 1.**
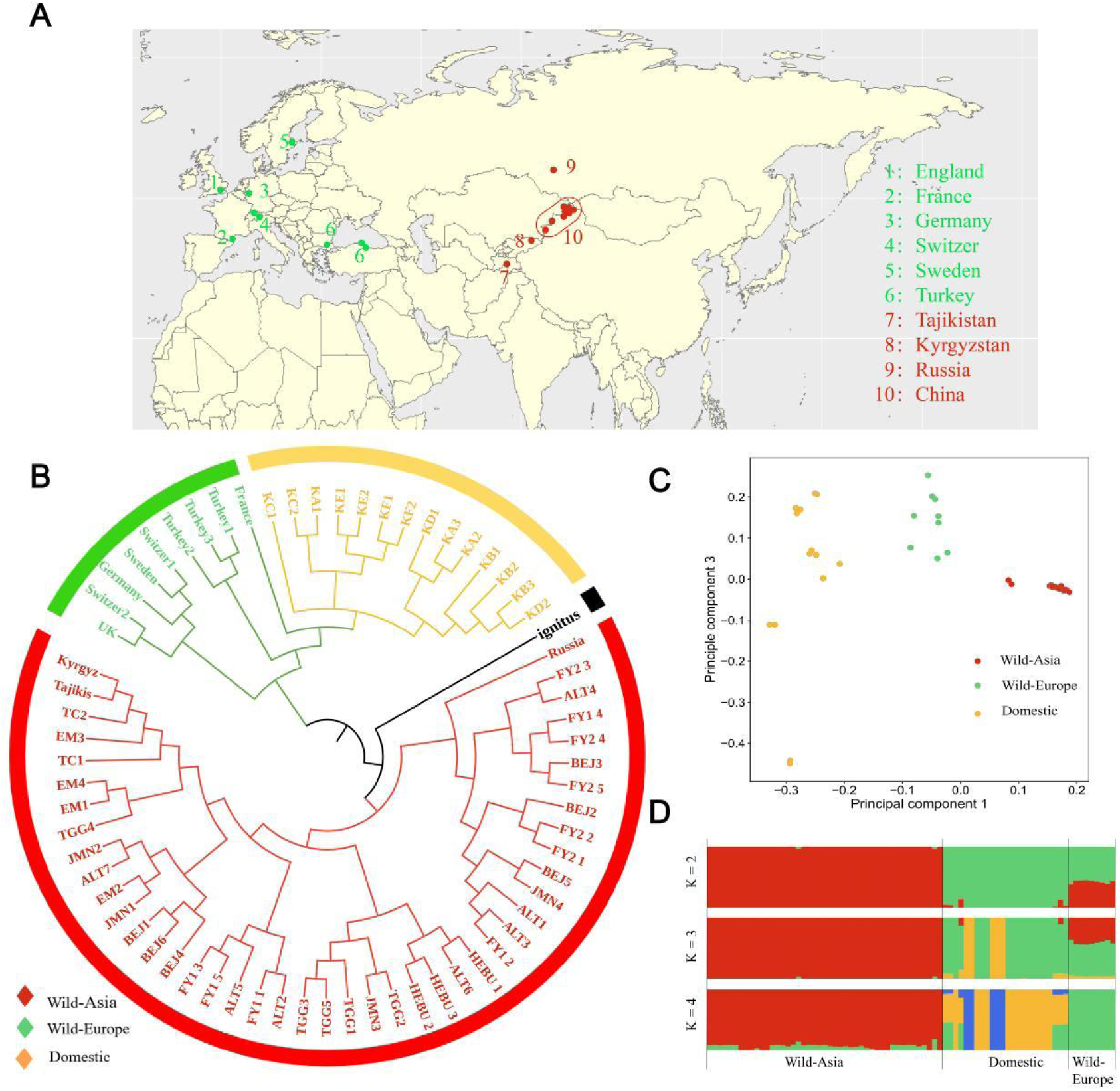
Sampling sites and population genomic analyses of *Bombus terrestris* individuals sampled from across Asia and Europe. (A) Geographic locations of the 53 wild-sampled *B. terrestris* individuals (red: collection sites in Asia, green: collection sites in Europe). (B) Maximum likelihood phylogenetic tree based on SNPs using *B. ignitus* as the outgroup, (C) Principal component analysis (PCA) plot showing wild-Asian (red), wild-European (green), and domestic (yellow) samples on PC1 and PC3, and (D) STRUCTURE analysis results (K = 2 - 4) for 67 of the 77 wild and domestic *B. terrestris* individuals (10 domestic samples excluded based on kinship analysis results).

To identify selected genomic regions in the Asian population, we employed three metrics: 1) the ratio of genetic diversity between European and Asian populations (ln-ratio _πEurope/πAsia_); 2) the population differentiation coefficient *F*_ST_ between European and Asian populations; and 3) the Composite Likelihood Ratio (CLR) values in Asian the population. The ratio of genetic diversity and the population differentiation coefficient *F*_ST_ were calculated with vcftools (versions 4.2) (Danecek et al., 2011) using a sliding window approach, with a window size of 10 kb and a step size of 5 kb. CLR values were calculated with SweeD (Pavlidis et al., 2013) using a grid size of 1 kb. Based on results from the above two analyses, genetic diversity ratios (ln-ratio _πEurope/πAsia_), *F*_ST_ values, and CLR values were ranked to identify genomic regions falling within the top 5% for all these three metrics, which were deemed as regions putatively under selection in the Asian *B. terrestris* population.

### Functional prediction of focal genes

To predict the functions of genes under selection in the Asian population, we first identified the genomic locations of these genes using the genome annotation file (Annotation Release 102). Subsequently, we extracted their amino acid sequences and used InterProScan software (Jones et al., 2014) to assign Gene Ontology (GO) terms for each protein-coding gene. The results of the GO annotation were then subjected to enrichment analysis using the R Package TopGO (Alexa et al., 2024). In addition, we utilized the Kyoto Encyclopedia of Genes and Genomes (KEGG) Orthology Based AnnotationSystem (KOBAS) (Xie et al., 2011) to perform KEGG pathway annotation and enrichment analysis for these genes. Finally, we performed BLAST searches using the amino acid sequences of these genes against the fruit fly database, FlyBase (Marygold et al., 2016), to identify homologous genes in *Drosophila*. Through this approach we could infer potential functions of these *B. terrestris* genes based on their counterparts in fruit flies.

## Results

### Genome resequencing of buff-tailed bumblebees collected from Europe and Asia

A total of 77 *B. terrestris* samples were collected from Asia and Europe for this analysis (Fig. 1A). Specifically, 41 wild samples were collected from 9 locations within Xinjiang province, China, with an average distance of 254 kilometers between collection sites. 12 additional wild samples were obtained from the Natural History Museum London (https://www.nhm.ac.uk/), which were specimens collected from 6 countries in Europe (France, Germany, Sweden, Switzerland, Turkey, and England) and 3 Asian regions (Tajikistan, Kyrgyzstan, and Russia (82.93N 55.01E)) (Table S1). The identity of all these wild samples was confirmed by DNA barcoding (Hebert et al. 2004). In addition to the wild samples, 24 commercial (domesticated) *B. terrestris* samples were included in this study, which were bought from *KOPPERT* - a company based in the Netherlands.

We performed whole-genome resequencing for these samples and achieved a mean depth of coverage of 30.80× and mean breadth of coverage of 99.37% (Table S2). Notably, specimens from the Natural History Museum London yielded an average depth of 20.99× (ranging from 11.3× to 33.74×) and a mean breadth of coverage of 98.94% (ranging from 96.9% to 99.54%) (Table S3). The Genome Analysis Toolkit (Mckenna et al., 2010) software was used to call variants for these 77 buff-tailed bumblebees, and a total of 4,827,687 high-quality single nucleotide polymorphisms (SNPs) and 615,899 high-quality insertion-deletions (INDELs) were obtained. Based on the obtained SNPs, we estimated kinship coefficients for these bumblebees and found that 10 commercial samples were full-or-half-sisters to other individuals (1st degree sibling: 0.177-0.354; Manichaikul et al., 2010). Therefore, we excluded these 10 samples and a total of 67 buff-tailed bumblebees were retained for downstream analysis.

### Distinct genetic differentiation exists between Asian and European buff-tailed **bumblebees**

To understand the genome-wide relationships and divergence between different *B. terrestris* samples, we constructed the maximum likelihood (ML) tree using these 67 resequenced individuals, using *Bombus ignitus* (belonging to subgenus *Bombus*) as the outgroup (Fig. 1B, Fig. S1).

The phylogenetic tree based on genome-wide SNPs revealed that the 67 *B. terrestris* individuals could be divided into two groups: 1) the first group, named as Wild-Asia, consisted of 44 wild *B. terrestris* samples collected from China, Tajikistan, Kyrgyzstan, and Russia; 2) the second group comprised 9 wild samples collected from France, Germany, Sweden, Switzerland, Turkey, and England (named as Wild-Europe) and 14 domesticated samples (named as Domestic) (Fig. 1B). In addition, the close relationship observed between the Domestic and Wild-Europe *B. terrestris* samples (Fig. 1B) was consistent with the fact that the commercial *B. terrestris* colonies have been domesticated from wild *B. terrestris* in Europe (Velthuis and Van Doorn 2006). The relationships inferred from the phylogenetic analysis were supported by Bayesian clustering analysis using principal component analysis (PCA) (Fig. 1C) and ADMIXTURE analysis (Fig. 1D). There was a clear genetic differentiation between *B. terrestris* samples originating from Asia and those from Europe.

To quantify the genetic differentiation between Asian and European *B. terrestris*, we calculated pairwise *F*_ST_ values. The results showed that the pairwise *F*_ST_ value between Wild-Asia and Wild-Europe was 0.076404 (Table. 1), indicating that there is moderate genetic differentiation between them (Spitze, 1993).

### Buff-tailed bumblebees in Asia dispersed from Europe

We calculated genetic diversity summary statistics and the results revealed that the nucleotide diversity (π) value for the Wild-Europe group (π = 0.002906) was higher than that of the Wild-Asia group (π = 0.002276), suggesting that buff-tailed bumblebee populations in Europe are older than those of Asia (Table. 1). In addition, Tajima’s D test [Tajima’s D (Wild-Europe) = -0.027101, Tajima’s D (Wild-Asia) = 1.100971] (Table 1) suggested that buff-tailed bumblebees in Europe have a larger population size than in Asia.

**Table 1.** Population genetic diversity summary statistics

| Population | Pi | Tajima's D | $F_{ST}$ | |
| --- | --- | --- | --- | --- |
|  |  |  | Wild-Asia | Domestic |
| Wild-Europe | 0.003117 | -0.02952 | 0.076122 | 0.060032 |
| Wild-Asia | 0.001877 | 1.8451 | - | 0.13277 |

To test the possibility that populations of Asian buff-tailed bumblebees were impacted by the invasion of commercial *B. terrestris* colonies (imported from Europe), we assessed gene flow between each population, specifically between the commercial samples and the samples collected from Xinjiang province. Firstly, we used TreeMix (Pickrell and Pritchard 2012) to infer the migration history among populations (Wild-Asia, Wild-Europe, Domestic) by using the joint allele frequencies among all populations in our study. TreeMix employs a maximum likelihood approach to model the genetic relationships among populations and detect admixture events. Our analysis did not reveal any gene flow between Domestic samples and Wild-Asia samples (Fig. 2); gene flow was found between Domestic samples and Wild-Europe samples, which is expected as the former was domesticated from the latter; gene flow was also found between Tajikistan and China-Xinjiang samples, which is expected as they are geographically adjacent (Fig. 2A; Fig. 2B; Fig. 1A). To check the above gene flow analysis between populations, we used the ABBA-BABA gene flow test. Results confirmed that no gene flow was observed between Wild-Asia (Xinjiang) samples and Domestic samples (Fig. 2C, 2D).

**Fig 2.**
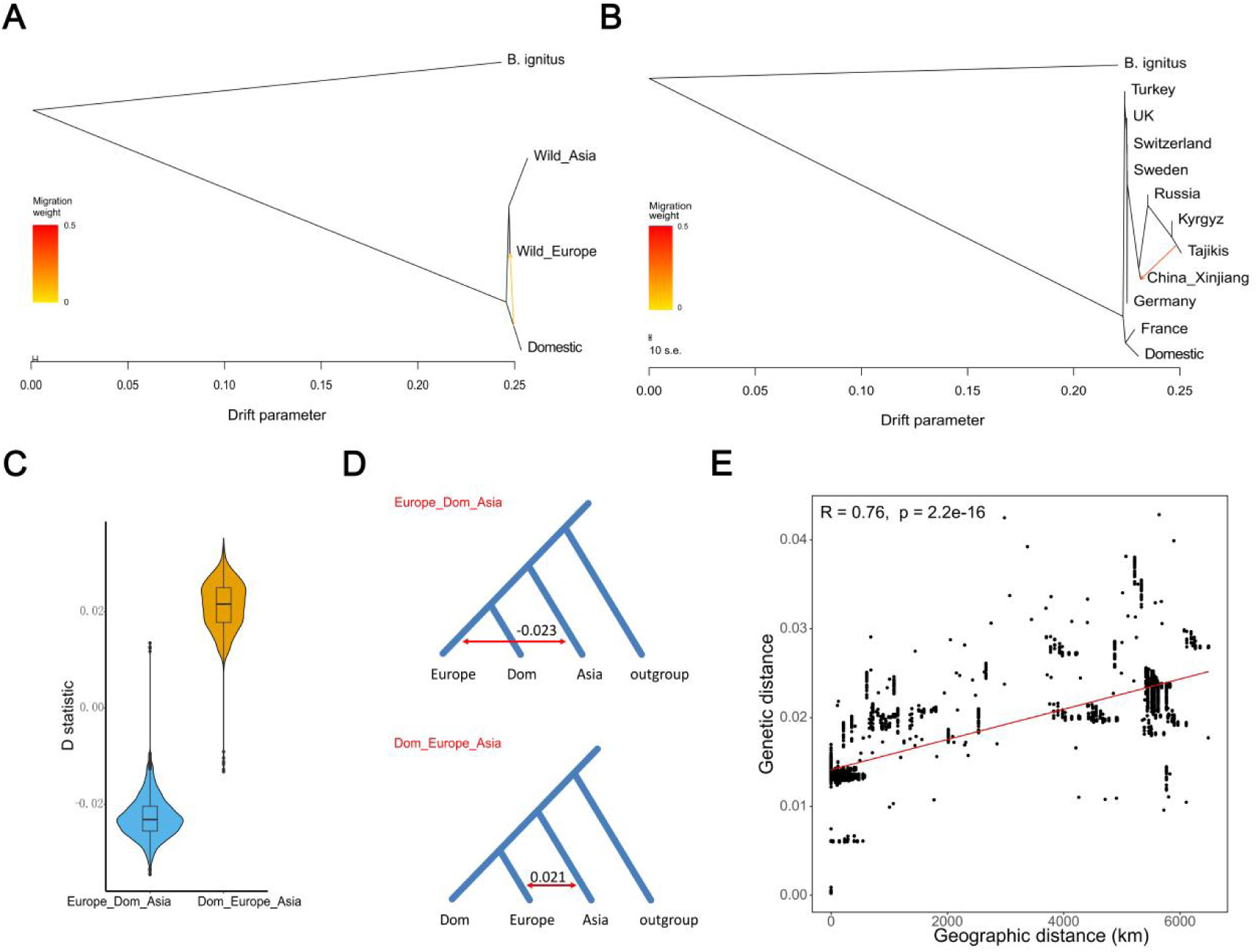
Gene flow analysis for *B. terrestris* samples collected from different geographic locations. (A) TreeMix gene flow analysis between the three main *B. terrestris* groups. (B) TreeMix gene flow analysis between different collection sites. (C, D) Results from the ABBA-BABA gene flow test. The topology of Europe_Dom_Asia is (((Wild-Europe, Domestic), Wild-Asia), outgroup), and Dom_Europe_Asia is (((Domestic, Wild-Europe), Wild-Asia), outgroup). (E) Correlation analysis of genetic and geographic distances of *B. terrestris* samples.

Finally, we calculated the geographic and genetic distances between wild *B. terrestris* samples in Europe and Asia. We found that the genetic distance between these samples follows a strict isolation-by-distance model (R = 0.76, p < 2.2e-16) and a correlation between genetic distance and geographic distance was found among individuals (Fig. 2E), suggesting that dispersal could explain the current genetic distances between wild *B. terrestris* samples in Europe and Asia. Taken together, all these findings suggest that buff-tailed bumblebees in Asia dispersed from ancestral populations in Europe.

### The demographic history of buff-tailed bumblebees

Pairwise Sequentially Markovian Coalescent (PSMC) model (Li and Durbin 2011) analysis was employed to infer the historical effective population size (*Ne*) of buff-tailed bumblebees. Because this analysis has a high requirement for sequencing coverage, we used the 14 commercial *B. terrestris* samples to represent the European population and the 42 wild *B. terrestris* samples collected from Xinjiang province, China to represent the Asian population. The results (Fig. 3) indicated that buff-tailed bumblebees in Asia and Europe exhibited similar and relatively stable *Ne* until ∼ 0.3 million years (Myr) ago. The population sizes of buff-tailed bumblebees in these two regions began to diverge at ∼ 0.3 Myr ago, approximately at the onset of the Penultimate Glaciation (Zheng et al. 2002). After this, both buff-tailed bumblebee populations exhibited an increasing *Ne* and *B. terrestris* in Europe showed a consistently higher *Ne* than that of Asia.

**Fig 3.**
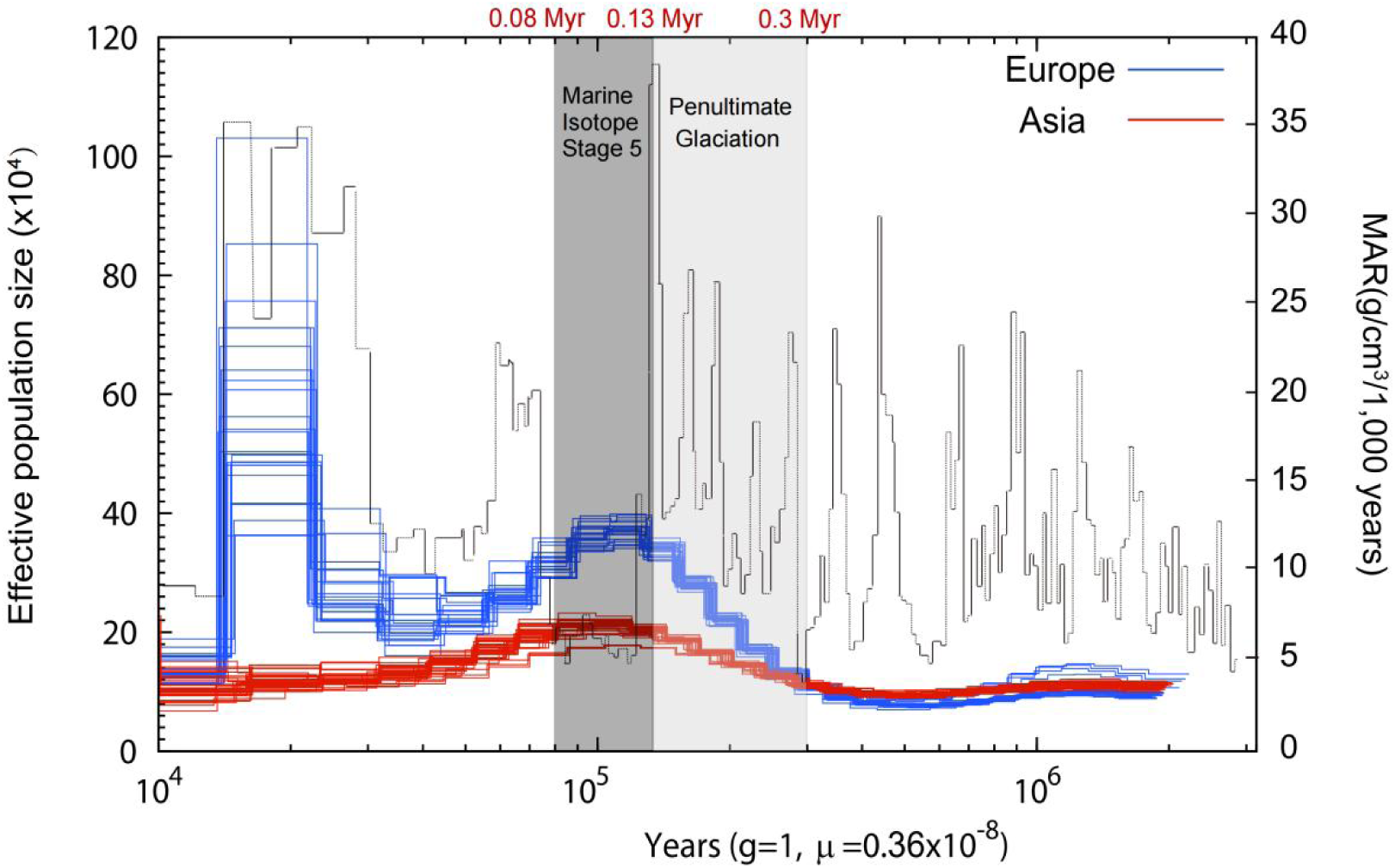
Demographic history of Asian and European *B. terrestris* populations inferred by software PSMC (g=1,µ=0.36×10-8). The gray shading marks the Penultimate Glaciation (PG; 0.13 – 0.3 Myr). The gray line represents the mass accumulation rate (MAR) of the Loess Plateau: high MAR values indicating cold and dry conditions, while low MAR values indicating hot and wet conditions.

The gray line in Figure 3 represents the mass accumulation rate (MAR) of the Loess Plateau (Sun and An 2005), which serves as an indicator of climatic conditions, with high MAR values indicating cold and dry periods and low MAR values indicating hot and wet periods. Comparing the climate and effective population size estimates over time, we observed two main trends: 1) the increase in *Ne* between 0.13 Myr ago and 0.3 Myr ago coincided with the Penultimate Glaciation period (0.13–0.3 million years ago) (Zheng et al. 2002), when the climate became cold and dry; 2) During Marine Isotope Stage 5 (MIS5, 0.08 Myr to 0.13 Myr ago), a period characterized by hot and wet climate according to the MAR estimates, we observed a decrease in *Ne* for *B. terrestris* both in Europe and Asia. Therefore, we infer from the demographic analysis that changes in *Ne* of buff-tailed bumblebees are associated with historical climate changes, with an increasing *Ne* during cold periods and a declining *Ne* during hot periods.

### Buff-tailed bumblebees in Asia and Europe differ in morphology and physiology

To understand if some environmental factors are significantly different between the two habitats (Europe and Asia) of buff-tailed bumblebees, we collated and compared the environmental factors of their habitats. Specifically, environmental data on temperature, elevation, solar radiation and precipitation were extracted from the global natural environment database (WorldClim) for the 24 *B. terrestris* collection sites in Europe and Asia. After comparison, we found several environmental factors were significantly different between the two habitats of *B. terrestris* (Fig. 4). Both the Mean and Minimum Temperatures of Asian habitats were significantly lower than that of European habitats (p_Ann-Mean-Temp_=0.02942; p_Mean-Temp-Cold_=0.02942; p_Min-Temp_=0.009444). Additionally, the Elevation and Sun Radiation in Asian habitats were significantly higher than in European habitats (p_Elevation_=0.007926; p_Sun-Radiation_=0.004594). Moreover, the Precipitation in the Asian habitats was significantly lower than in European habitats (p_Precipitation_=0.001733).

**Fig 4.**
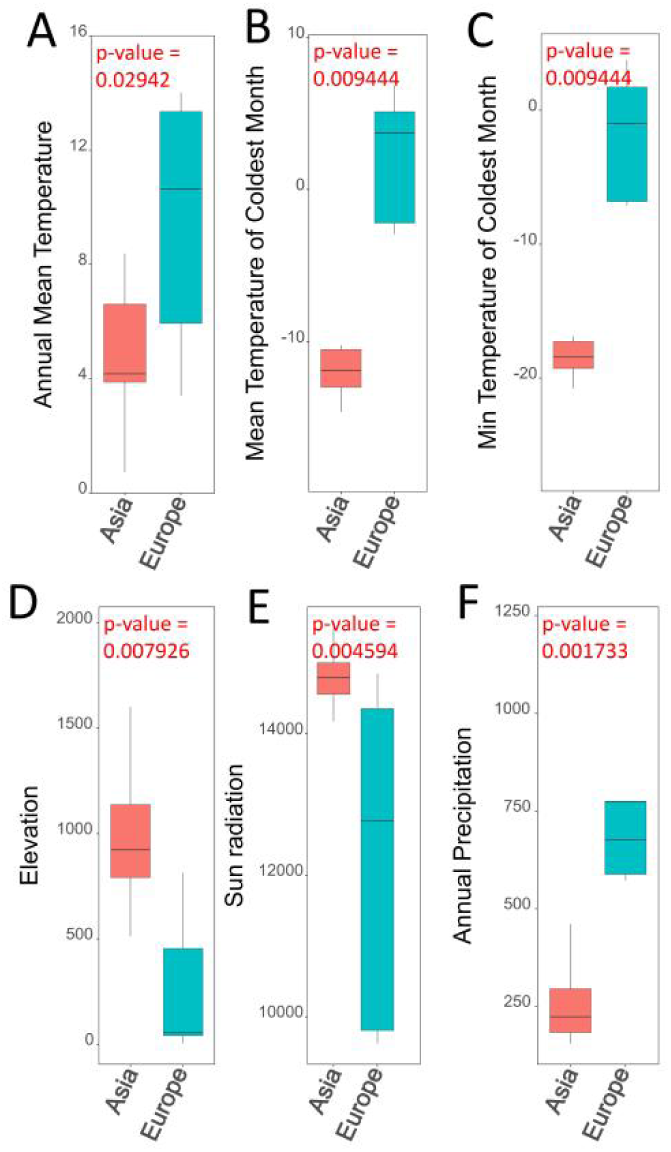
The comparison of climate factors between the habitats of Asian and European *B. terrestris*, which include: (A) annual mean temperature, (B) mean temperature of coldest month, (C) minimum temperature of coldest month, (D) elevation, (E) solar radiation, and (F) annual precipitation. P-values from Wilcoxon rank-sum tests are indicated for each comparison.

To understand if buff-tailed bumblebees in the two habitats exhibit any phenotypic differences, we measured and compared a selection of morphological characters. We randomly selected Asian and European buff-tailed bumblebees from different colonies and measured a set of features, including proboscis length, forewing length (FL), forewing breadth (FB), cubital vein A, cubital vein B, wing vein angles (A4, B4, D7, E9, G18, J10, J16, K19, L13, N23, O26, new angle), hooks on the right hind wing, femur length (Fe) of the right hind leg, tibia length (Ti) of the right hind leg, metabasitarsus length (ML), metabasitarsus breadth (MT), length of the third abdominal sternum (S3), breadth of the third abdominal sternum, breadth of the fourth abdominal sternum, length of the genitalia, breadth of the genitalia (Table S4). Results indicated that buff-tailed bumblebees in Asia had a shorter proboscis length compared to their European counterparts (Fig. 5B, 5E). The proboscis lengths of Asian queens (n=10), workers (n=30), and males (n=15) were 9.75±0.44, 6.75±1.11, and 8.33±1.44, respectively, while these lengths for European queens (n=10), workers (n=30), and males (n=15) were 11.74 ± 0.85, 9.24 ± 0.81, and 10.03 ± 0.72, respectively. We also measured the hair density of the buff-tailed bumblebees, which refers to the number of small pores within a 0.04-centimeter diameter area on a surface, and found Asian samples exhibited a higher hair density (Fig. 5A, 5C, 5D, 5G). The hair densities on the face and mesothorax of Asian bumblebees (n=30) were 775.1±39.6 and 863.4 ± 50.2, respectively, whereas those lengths for European bumblebees (n=30) were 675.97±78.75 and 594.5±53.56, respectively.

**Fig 5.**
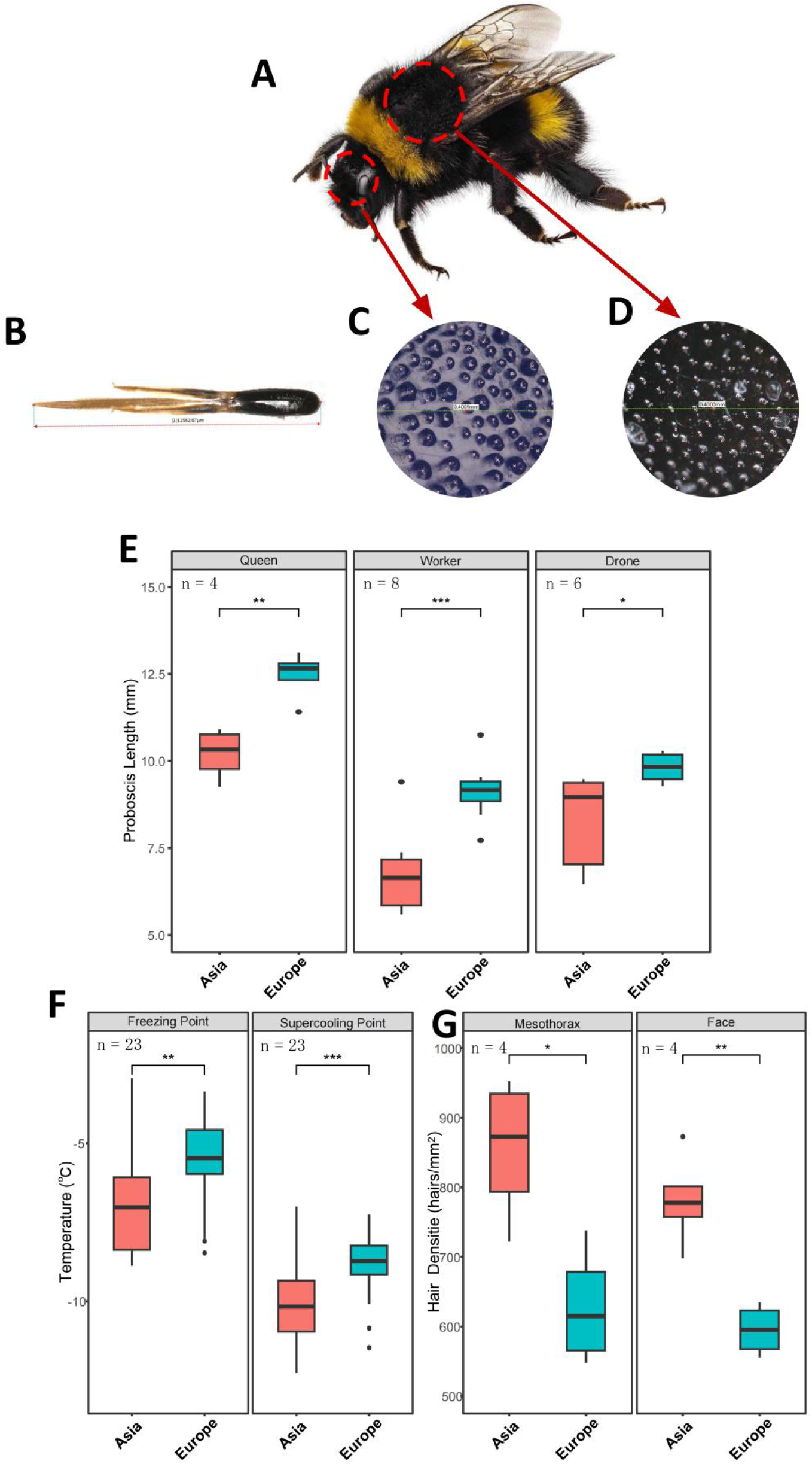
The morphological and physiological differences detected between Asian and European *B. terrestris*. The overall appearance (A), proboscis (B), hair density on the face (C), and mesothorax (D), of Asian *B. terrestris*. (E) Comparative analysis of proboscis lengths for the queens, workers, and drones of Asian and European *B. terrestris*. (F) The freezing points and supercooling points of Asian and European *B. terrestris*. (G) The hair density on the face and mesothorax of Asian and European *B. terrestris*. P-values are indicated with *: p<0.05, **: p<0.01, ***: p<0.005.

**Fig 6.**
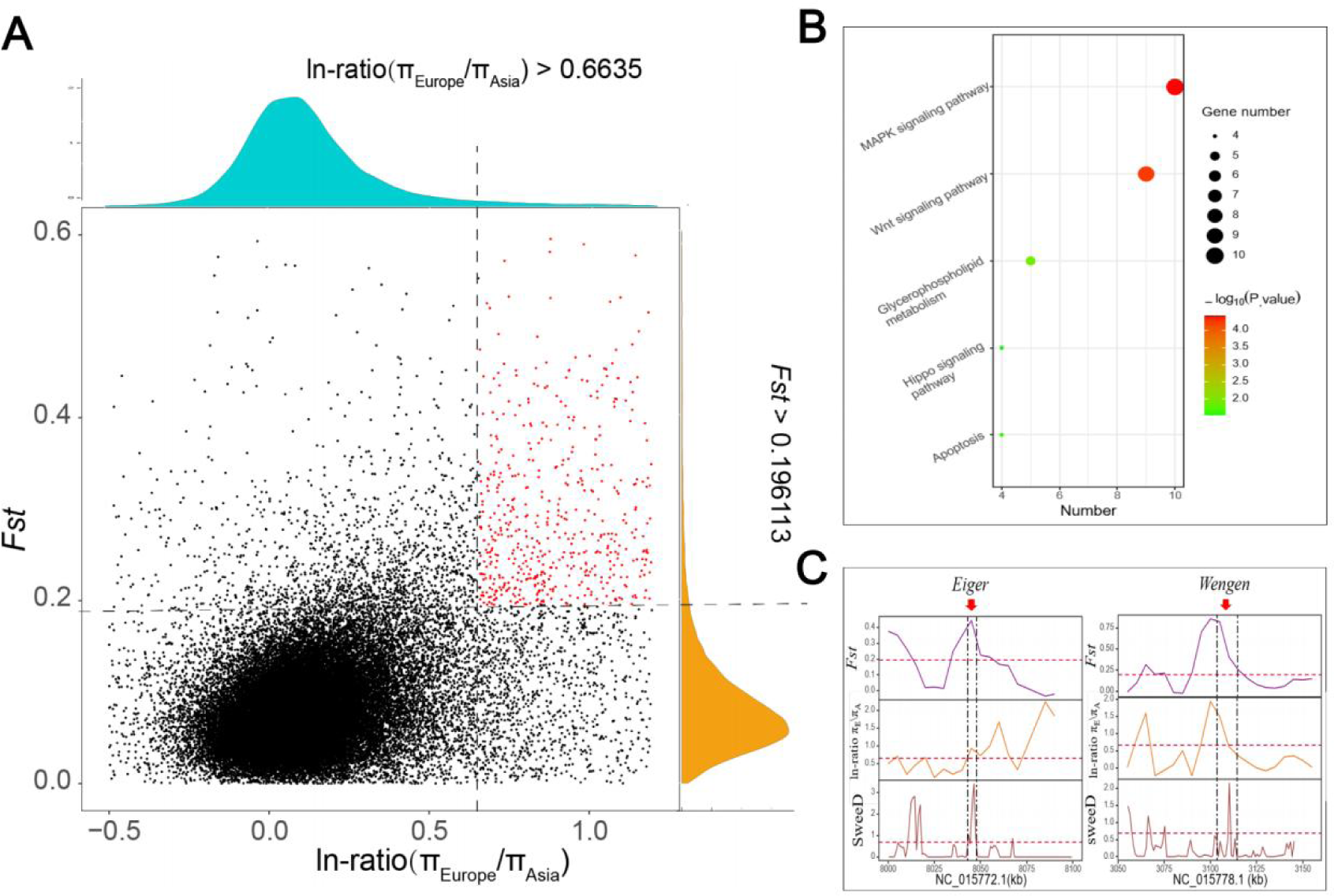
Genomic regions and genes showing evidence of positive selection in Asian *B. terrestris*. (A) Distribution of ln-ratio πEurope/πAsia and fixation statistics *F*ST of 10 kb windows with 5 kb steps. Red dots represent windows fulfilling the selected regions requirement (based on the top 5% for ln-ratio πEurope/πAsia (>0.6635) and fixation statistics *F*ST (>0.196113)). (B) The plot shows the Gene Ontology (GO) categorization of identified genes in significant pathways (p <0.05). The color of the circle represents the − log P.value. The size of the circle represents the number of genes under selection in the enriched pathway. (C) Two focal genes – *Eiger* and *Wengen* - showing selection signals in Asian *B. terrestris*. ln-ratio πEurope/πAsia, *F*ST, and SweeD (> 0.686) values are plotted using a 10kb sliding window.

**Fig 7.**
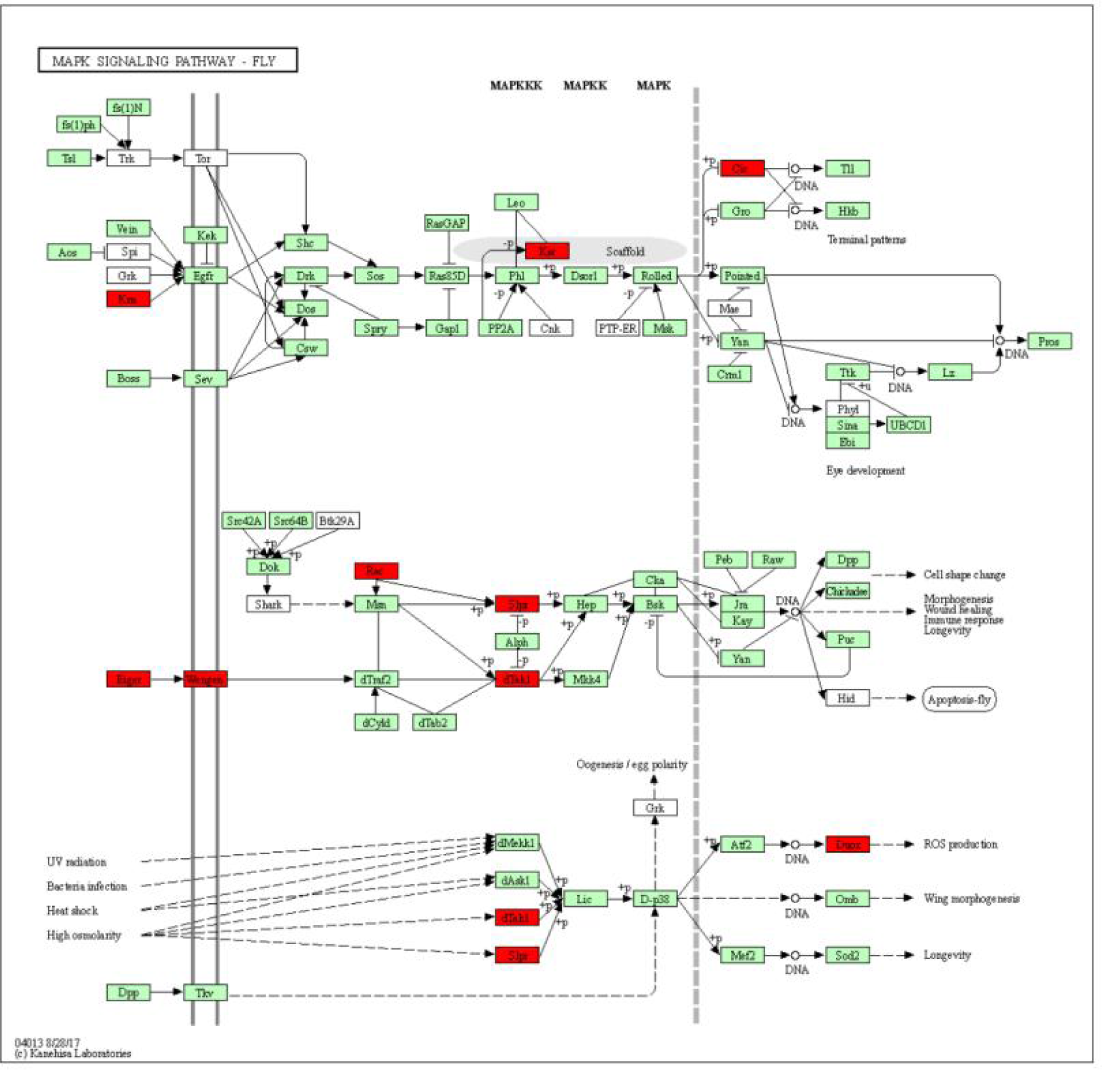
The Kyoto Encyclopedia of Genes and Genomes (KEGG) pathway for MAPK signaling. Genes in the MAPK signaling pathway that exhibited patterns of positive selection in Asian *B. terrestris* are highlighted in red.

The ambient temperature in the Asian habitats of buff-tailed bumblebees was lower than that of European buff-tailed bumblebees (Fig. 5F), so Asian *B. terrestris* may have an enhanced capacity for cold tolerance. As the supercooling point and freezing point were important parameters related to the survival of insects in cold environments, we conducted measurements of the supercooling point and freezing point for buff-tailed bumblebees in Asia and Europe. Results suggested that, as expected, the supercooling point and freezing point of Asian buff-tailed bumblebees were significantly lower than those of their European counterparts (Fig. 5F). The supercooling point of *B. terrestris* in Asia was -10.17±1.26 ℃ and the freezing point was -7.02 ± 1.5 ℃. In contrast, the supercooling point of European *B. terrestris* was -8.85±1.04 ℃, with a freezing point of -5.58±1.34 ℃.

### Genomic signatures of local adaptation for buff-tailed bumblebees in Asia

The phenotypic and physiological differences observed in Asian buff-tailed bumblebees indicate that Asian *B. terrestris* have undergone local adaptation to their Asian habitats. To identify genomic regions that experienced positive selection in Asian *B. terrestris*, we performed whole-genome genetic differentiation analysis between Asian samples and European samples. We used the Sliding-Window method, with a window size of 10 kb and a step size of 5 kb, based on the top 5% for ln-ratio πEurope/πAsia (>0.6635) and fixation statistics *F*_ST_ (>0.196113) (Axelsson et al., 2013; Qiu et al., 2015). Our analysis yielded a total of 435 candidate genomic regions that were under positive selection encompassing ∼ 6.64 Mb (2.64%) of the *B. terrestris* reference genome, with an average size of 15.2 Kb. Within these regions, there were 712 predicted protein-coding genes. To further refine this result, we employed the Composite Likelihood Ratio (CLR) test to identify genomic regions that were under selection. Candidate regions were extracted if they fell within the top 5% of Z-transformed values (>0.686) and they were identified as candidate regions under selection by the above ln-ratioπ and *F*_ST_ analysis. Applying these filters resulted in a total of 181 genomic regions being identified as under selection, within which there are 331 protein-coding genes (Table S7).

## Discussion

The buff-tailed bumblebee (*B. terrestris*) is an important pollinator of wild plants and is one of the most widely used pollinator species for greenhouse crop pollination (Velthuis and Van Doorn 2006; Goulson, 2010). Intensive studies have been performed to understand its population structure, genetic differentiation and adaptation using samples from European and nearby Mediterranean and Atlantic Island populations (Moreira et al., 2015; Kraus et al., 2009; Lecocq et al., 2013; Estoup et al., 1996; Widmer et al. 1998; Silva et al., 2020; Colgan et al., 2022). However, less was known about *B. terrestris* populations in Asia, which represent the eastern edge of its global natural distribution (Williams et al. 2012). In this study, we collected wild *B. terrestris* samples from 11 geographic populations in Asia, as well as wild samples from Europe and domesticated samples from a commercial company in Europe. After whole-genome resequencing of these samples, we investigated their population genetic structure, demographic history and local adaptation.

Our results indicated that buff-tailed bumblebees could be divided into two groups: while samples originating from Asia were clustered into one group, samples originating from Europe were clustered into another group (Fig. 1). That is, previous studies on buff-tailed bumblebees largely missed the diversity represented by Asian populations. In addition, although *B. terrestris* in Asia most likely dispersed from European populations (Fig. 2; Fig. 3), distinct genetic differentiation was detected between the two groups (Table 1), with a *F*_ST_ value of 0.076404, indicating a moderate genetic differentiation (Spitze, 1993). Moreover, buff-tailed bumblebees in Asia and in Europe differ in morphology and physiology. For example, compared to their European counterparts, buff-tailed bumblebees collected in Asia (more specifically, Xinjiang province, China), had a shorter proboscis length, a higher hair density and a lower supercooling and freezing point (Fig. 5; Table S4). Taken together, considering the moderate genetic differentiation and distinct morphological and physiological differences,, buff-tailed bumblebees in Asia represent a distinct new genetic resource.

### Description of *Bombus terrestris* in Asia

*Distribution*: The species is mainly distributed in Asia in Iran, Russia, and the mountains of Central Asia and the northern Xinjiang region of China - areas of the Tianshan Mountains and the Altai Mountains, but also including western Mongolia.

*Color Pattern*: Consists of alternating bands of black and yellow on its body. The fore margin of thorax is covered in dense yellow hairs; the hind margin of abdomen is covered in white hairs; black elsewhere; corbicular cuticle and setae black.

*Morphology*: Morphological parameters of *B. terrestris* in Asia, including proboscis length, forewing length (FL), forewing width (FB), cubital vein A, cubital vein B, wing vein angles (A4, B4, D7, E9, G18, J10, J16, K19, L13, N23, O26, new angle), hooks on the right hind wing, femur length (Fe) of the right hind leg, tibia length (Ti) of the right hind leg, metabasitarsus length (ML), metabasitarsus breadth (MT), length of the third abdominal sternum (S3), width of the third abdominal sternum, width of the fourth abdominal sternum, length of the genitalia, breadth of the genitalia. Compared to the European bumblebees, *B. terrestris* in Asia have shorter proboscis length, higher hair density. All the morphometric characters are presented in supplementary Table S4, Supplementary Material online.

### The local adaptation of buff-tailed bumblebees in Asia

Local adaptation is an important ability required for organisms to survive in changing environments (Fuse 2017). The dispersal of *B. terrestris* from Europe to Asia has encountered different environmental conditions, including those of temperature, elevation, and precipitation (Fig. 4). Correspondingly, *B. terrestris* in Asia exhibited altered phenotypes after occupying new environments, such as changes in proboscis length, increased hair density, and lower supercooling point (Fig. 5). It has been reported that plants at higher altitudes have smaller overall sizes of flowers (Ansari et al. 2010; Zhu et al. 2010; Halbritter et al. 2018; Kiełtyk et al. 2021). Therefore, flower sizes in *B. terrestris’* Asian habitats, which have a higher elevation (Fig. 4), should be smaller than in European habitats. Therefore, the shorter proboscis observed in *B. terrestris* in Asia could represent an adaptive trait for them to collect food. Both mean and minimum temperatures of *B. terrestris’* habitat in Asia were significantly lower than in Europe (Fig. 4). Bumblebees primarily rely on their body hair to withstand low-temperature challenges (Heinrich et al. 1974), so the increased hair density in *B. terrestris* in Asia could provide them with enhanced ability to withstand cold temperatures. The phenomenon of supercooling refers to the ability of insects to remain in a liquid state even when the temperature is below their freezing point, which is widespread among insects and is crucial for their survival in cold environments (Renault et al. 2002; Lombardo et al. 2017). A comparison between *B. terrestris* in Asia and the European *B. terrestris* revealed that the former had a lower supercooling point (Fig. 5), suggesting that *B. terrestris* in Asia is adapted to low temperatures.

Genes related with local adaptation will show positive selection signals within the genome. Here we employed a method that has been used in several organisms, such as yaks (Qiu et al., 2015a), cows (Chen et al., 2018b), silkworms (Xia et al., 2009), pigs (Rubin et al., 2012), soybeans (Zhou et al., 2015), and rice (Huang et al., 2012), to identify genes under positive selection in *B. terrestris* in Asia. Our results indicate that the MAPK pathway and glycerophospholipid (GPL) metabolic pathway in *B. terrestris* in Asia had undergone selection (Table S6). Specifically, we observed that *Eiger* and *Wengen* in these two pathways have been subjected to positive selection (Table S7). Previous research has shown that *Eiger*, a homolog of tumor necrosis factor in fruit flies, is released in epidermal cells undergoing apoptosis in response to intense UV radiation; *Wengen*, on the other hand, acts as the receptor for *Eiger* on sensory neurons, mediating the transmission of adaptive signals that enable the organism to cope with UV radiation (Agrawal et al., 2016). Therefore, it is likely that *Eiger* and *Wengen* contributed to the adaptation of *B. terrestris* in Asia to the high-elevation and UV-intensive environments in their habitats. Additionally, studies have demonstrated that *Eiger* can modulate insulin-mediated nutritional responses in fruit flies, enhancing their adaptability to changing nutritional environments (Agrawal et al., 2016). Considering the habitat characteristics of *B. terrestris* in Asia, such as low precipitation, high elevation, and limited vegetation leading to reduced food availability (Fig. 4), *Eiger* may play a role in enhancing Asian *B. terrestris’* adaptability to their nutritional environments. Genes in the glycerophospholipid (GPL) metabolic pathway were also found under positive selection (Table S6). Previous studies demonstrated the association between the glycerophospholipid (GPL) metabolism pathway and low-temperature adaptation in the hemipteran insect *Pyrrhocoris apterus* (Han et al., 1995; Tomcala et al., 2006). Also, it was reported that glycerophospholipids (GPL) play a crucial role in regulating the supercooling point in insects (Tomcala et al., 2006). Therefore, genes in the glycerophospholipid (GPL) metabolic pathway that showed selection signals likely enhanced the cold tolerance of *B. terrestris* in Asia.

## Conclusion

Through collecting and re-sequencing buff-tailed bumblebees from their Asian habitats, which were less well represented in previous studies, we performed the first multi-locus population genetic structure analysis of this important pollinator. Our results revealed that buff-tailed bumblebees in Asia dispersed from Europe and accumulated a moderate genetic differentiation with their European counterparts. Buff-tailed bumblebees in Asia adapted to their habitats morphologically and physiologically and we identified genes that were likely involved in their adaptation to high UV radiation, low temperature, and low precipitation of their habitats. The new genomic resources and insights revealed through this study will be valuable for the conservation and improved management of this important pollinator.

## Author contributions

C.S. and Y.Z. conceived and designed the study; P.H.W., L.S., C.S. and Y.Z. collected bee samples; L.S., D.L., Y.C.L., R. W., X. D., S.Z., H. F., X.Z., H.S. and Q.W. were involved in methodology; L.S., Y.C.L. collaborated on script compilation for data analysis; L.S. and C.S. drafted the initial manuscript; R.M.W., P.H.W. and Y.Z. gave guidance in the writing; all authors participated in reviewing and editing the manuscript.

## Supporting information

Supplemental Table 1 Summary of wild samples information

Supplemental Table 2 Quality control data

Supplemental Table 3 Resequencing Mapping data

Supplemental Table 4 Morphological parameters of Bombus terrestris from Asia and Europe

Supplemental Table 5 GO enrichment analysis of Asia samples selected genes

Supplemental Table 6 KEGG enrichment analysis of Asia samples selected genes

Supplemental Table 7 Asia samples selected genes

## Acknowledgements

This work was supported by the National Natural Science Foundation of China (Grant No. 32270445 and 31971397), Swiss National Science Foundation (SNSF) (Grant numbers 202669 and 196125 to RMW), Shandong Provincial Natural Science Foundation (ZR2021YQ21, ZR2021QC218); Shandong Provincial Agriculture Research System (SDAIT-24); Taishan Scholars program of Shandong Province (tsqn202312293); Shandong Provincial Key R&D Program (2023LZGC017).

This work was also supported by the Shandong Academy of Agricultural Sciences, Institute of Plant Protection, Shandong Key Laboratory for Green Prevention and Control of Agricultural Pests, Shandong Provincial Engineering Research Center for Development and Utilization of Beneficial Insects.

## Data and Code Availability Statement

All data and code will be made available upon acceptance.

## SUPPLEMENTARY MATERIAL

**Table S1.** Summary of wild samples information

| Sample | Site_country | Site_province | Site_name | Collect_date | Longitude | Latitude |
| --- | --- | --- | --- | --- | --- | --- |
| ALT1 | China | Xinjiang | Altay | 2019.6 | 88.1411°E | 47.8522°N |
| ALT2 | China | Xinjiang | Altay | 2019.6 | 88.1411°E | 47.8522°N |
| ALT3 | China | Xinjiang | Altay | 2019.6 | 88.1411°E | 47.8522°N |
| ALT4 | China | Xinjiang | Altay | 2019.6 | 88.1411°E | 47.8522°N |
| ALT5 | China | Xinjiang | Altay | 2019.6 | 88.1411°E | 47.8522°N |
| ALT6 | China | Xinjiang | Buerjin | 2018.6 | 86.52°E | 47.42°N |
| ALT7 | China | Xinjiang | Buerjin | 2018.6 | 86.52°E | 47.42°N |
| BEJ1 | China | Xinjiang | Buerjin | 2019.6 | 86.52°E | 47.42°N |
| BEJ2 | China | Xinjiang | Buerjin | 2019.6 | 86.52°E | 47.42°N |
| BEJ3 | China | Xinjiang | Buerjin | 2019.6 | 86.52°E | 47.42°N |
| BEJ4 | China | Xinjiang | Buerjin | 2019.6 | 86.52°E | 47.42°N |
| BEJ5 | China | Xinjiang | Buerjin | 2019.6 | 86.52°E | 47.42°N |
| BEJ6 | China | Xinjiang | Buerjin | 2018.6 | 86.52°E | 47.42°N |
| EM1 | China | Xinjiang | Emin | 2019.6 | 83.6133°E | 46.5238°N |
| EM2 | China | Xinjiang | Emin | 2019.6 | 83.6133°E | 46.5238°N |
| EM3 | China | Xinjiang | Emin | 2019.6 | 83.6133°E | 46.5238°N |
| EM4 | China | Xinjiang | Emin | 2019.6 | 83.6133°E | 46.5238°N |
| FY1-1 | China | Xinjiang | Fuyun | 2019.6 | 89.535°E | 46.9811°N |
| FY1-2 | China | Xinjiang | Fuyun | 2019.6 | 89.535°E | 46.9811°N |
| FY1-3 | China | Xinjiang | Fuyun | 2019.6 | 89.535°E | 46.9811°N |
| FY1-4 | China | Xinjiang | Fuyun | 2019.6 | 89.535°E | 46.9811°N |
| FY1-5 | China | Xinjiang | Fuyun | 2019.6 | 89.535°E | 46.9811°N |
| FY2-1 | China | Xinjiang | Fuyun | 2019.6 | 89.8233°E | 47.2141°N |
| FY2-2 | China | Xinjiang | Fuyun | 2019.6 | 89.8233°E | 47.2141°N |
| FY2-3 | China | Xinjiang | Fuyun | 2019.6 | 89.8233°E | 47.2141°N |
| FY2-4 | China | Xinjiang | Fuyun | 2019.6 | 89.8233°E | 47.2141°N |
| FY2-5 | China | Xinjiang | Fuyun | 2019.6 | 89.8233°E | 47.2141°N |
| HEBU1 | China | Xinjiang | Hoboksar | 2019.6 | 85.7236°E | 46.7869°N |
| HEBU2 | China | Xinjiang | Hoboksar | 2019.6 | 85.7236°E | 46.7869°N |
| HEBU3 | China | Xinjiang | Hoboksar | 2019.6 | 85.7236°E | 46.7869°N |
| JMN1 | China | Xinjiang | Jimunai | 2019.6 | 85.8697°E | 47.4169°N |
| JMN2 | China | Xinjiang | Jimunai | 2019.6 | 85.8697°E | 47.4169°N |
| JMN3 | China | Xinjiang | Jimunai | 2019.6 | 85.8697°E | 47.4169°N |
| JMN4 | China | Xinjiang | Jimunai | 2019.6 | 85.8697°E | 47.4169°N |
| TC1 | China | Xinjiang | Tacheng | 2018.6 | 82.5189°E | 45.9499°N |
| TC2 | China | Xinjiang | Tacheng | 2018.6 | 82.5189°E | 45.9499°N |
| TGG1 | China | Xinjiang | Tiechanggou | 2019.6 | 84.4494°E | 46.1672°N |
| TGG2 | China | Xinjiang | Tiechanggou | 2019.6 | 84.4494°E | 46.1672°N |
| TGG3 | China | Xinjiang | Tiechanggou | 2019.6 | 84.4494°E | 46.1672°N |
| TGG4 | China | Xinjiang | Tiechanggou | 2019.6 | 84.4494°E | 46.1672°N |
| TGG5 | China | Xinjiang | Tiechanggou | 2019.6 | 84.4494°E | 46.1672°N |
| Fra | France | Perpignan | Perpignan | 2018.1 | 2.89541°E | 42.6976°N |
| Ger | Germany | NR-Westfalia | Muenster | 2011.4 | 6.953101°E | 50.9351°N |
| Kyr | Kyrgyzstan | Issyk-Kul | Barskoon | 2009.7 | 77.5°E | 42.5°N |
| Rus | Russia | Novosibirsk | Chany lake | 2006.5 | 82.9339°E | 55.0188°N |
| Swe | Sweden | Uppsala | Predikstolen | 2011.4 | 17.6388°E | 59.8588°N |
| Swi1 | Switzerland | Zuerich | Hedingen | 2011.4 | 8.55°E | 47.3667°N |
| Swi2 | Switzerland | Graubunden | San Bernardino | 2011.4 | 9.54972°E | 46.6497°N |
| Taj | Tajikistan | Kuhistoni Badakhshon | Shirg | 2009.5 | 71.2715°E | 38.3234°N |
| Tur1 | Turkey | Sinop | Alaçam-Dur ağan | 2010.5 | 35.1625°E | 42.0268°N |
| Tur2 | Turkey | Edirne | İbriktepe-Uzunköprü | 2010.5 | 26.5834°E | 41.6749°N |
| Tur3 | Turkey | Samsun | Alaçam-Dur ağan | 2010.5 | 36.3361°E | 41.2797°N |
| UK | UK | England | Shortlands | 2011.4 | 0.136439°W | 51.5073°N |

**Table S2.** Quality control data

| Sample | Raw Base(bp) | Clean Base(bp) | Effective Rate(%) | Error Rate(%) | Q20(%) | Q30(%) | GC Content(%) |
| --- | --- | --- | --- | --- | --- | --- | --- |
| ALT1 | 11204490000 | 11187918300 | 99.85 | 0.03 | 97.72 | 94.06 | 39.61 |
| ALT2 | 11115318900 | 11099028900 | 99.85 | 0.03 | 98.05 | 94.65 | 39.61 |
| ALT3 | 11735084700 | 11717913300 | 99.85 | 0.03 | 98.01 | 94.37 | 39.6 |
| ALT4 | 11775414600 | 11758554000 | 99.86 | 0.03 | 97.89 | 94.22 | 39.22 |
| ALT5 | 11232938700 | 11214195900 | 99.83 | 0.03 | 97.48 | 93.5 | 40.71 |
| ALT6 | 23198895000 | 22913028300 | 98.77 | 0.04 | 94.15 | 87.45 | 39.04 |
| ALT7 | 23425786800 | 23162517900 | 98.88 | 0.04 | 94.81 | 88.52 | 38.56 |
| BEJ1 | 18807973500 | 18740560200 | 99.64 | 0.03 | 96.41 | 91.58 | 38.34 |
| BEJ2 | 15412268100 | 15361805100 | 99.67 | 0.03 | 96.11 | 90.62 | 38.13 |
| BEJ3 | 17629497000 | 17567310000 | 99.65 | 0.03 | 96.66 | 91.97 | 38.23 |
| BEJ4 | 14801996400 | 14752334400 | 99.66 | 0.03 | 96.57 | 91.67 | 38.25 |
| BEJ5 | 18776677200 | 18711360000 | 99.65 | 0.03 | 96.13 | 91.13 | 38.92 |
| BEJ6 | 23495247000 | 23403879600 | 99.61 | 0.03 | 96.25 | 91.45 | 39.09 |
| EM1 | 11361333600 | 11344302600 | 99.85 | 0.03 | 97.83 | 94.18 | 40.6 |
| EM2 | 11335378500 | 11318334600 | 99.85 | 0.03 | 97.94 | 94.33 | 40.89 |
| EM3 | 11329419600 | 11313071100 | 99.86 | 0.03 | 97.88 | 94.18 | 39.71 |
| EM4 | 11083358100 | 11066329200 | 99.85 | 0.03 | 97.58 | 93.5 | 40.49 |
| Fra | 10276995564 | 10296814276 | 99.19 | 0.02 | 97.92 | 91.96 | 36.24 |
| FY1_1 | 11684118900 | 11666343000 | 99.85 | 0.03 | 97.59 | 93.54 | 39.94 |
| FY1_2 | 11458690500 | 11441256000 | 99.85 | 0.03 | 97.69 | 93.86 | 39.81 |
| FY1_3 | 11383016100 | 11367117900 | 99.86 | 0.03 | 98.05 | 94.41 | 39.19 |
| FY1_4 | 11297316300 | 11280502200 | 99.85 | 0.03 | 97.9 | 94.1 | 39.99 |
| FY1_5 | 11012565600 | 10996983000 | 99.86 | 0.03 | 98.17 | 94.65 | 39.01 |
| FY2_1 | 11535843300 | 11518626300 | 99.85 | 0.03 | 97.84 | 94 | 39.84 |
| FY2_2 | 11772026100 | 11755001700 | 99.86 | 0.03 | 97.98 | 94.43 | 39.51 |
| FY2_3 | 11252029200 | 11235295500 | 99.85 | 0.03 | 97.96 | 94.17 | 39.46 |
| FY2_4 | 11187132600 | 11170856100 | 99.85 | 0.03 | 98.14 | 94.57 | 39.43 |
| FY2_5 | 11267707200 | 11251611000 | 99.86 | 0.03 | 98.08 | 94.37 | 39.07 |
| Ger | 19210869156 | 19259124154 | 99.25 | 0.02 | 99.16 | 94.04 | 36.89 |
| HEBU_1 | 11825811300 | 11807268900 | 99.84 | 0.03 | 97.61 | 93.84 | 40.51 |
| HEBU_2 | 11267631600 | 11250558600 | 99.85 | 0.03 | 97.68 | 93.88 | 40.19 |
| HEBU_3 | 11960218800 | 11942794200 | 99.85 | 0.03 | 97.9 | 94.34 | 39.94 |
| JMN1 | 11800650000 | 11782890900 | 99.85 | 0.03 | 97.6 | 93.77 | 39.76 |
| JMN2 | 11720851800 | 11702907000 | 99.85 | 0.03 | 97.44 | 93.11 | 40.29 |
| JMN3 | 11955773400 | 11937270000 | 99.85 | 0.03 | 97.58 | 93.41 | 40.56 |
| JMN4 | 11885277900 | 11867463300 | 99.85 | 0.03 | 97.67 | 93.81 | 40.03 |
| KA1 | 21588184200 | 21485203500 | 99.52 | 0.03 | 95.92 | 90.17 | 37.87 |
| KA2 | 11280674700 | 11264512800 | 99.86 | 0.03 | 97.42 | 93.13 | 39.98 |
| KA3 | 11446620600 | 11430573300 | 99.86 | 0.03 | 97.45 | 93.22 | 39.85 |
| KB1 | 11750325000 | 11732830200 | 99.85 | 0.03 | 97.42 | 93.22 | 41.18 |
| KB2 | 11186355600 | 11170710600 | 99.86 | 0.03 | 97.51 | 93.16 | 40.15 |
| KB3 | 14625807300 | 14542490700 | 99.43 | 0.04 | 95.74 | 89.97 | 37.69 |
| KC1 | 11661643500 | 11645530800 | 99.86 | 0.03 | 97.72 | 93.55 | 39.65 |
| KC2 | 14553318900 | 14468098800 | 99.41 | 0.03 | 95.87 | 90.24 | 37.85 |
| KD1 | 14658367800 | 14577426600 | 99.45 | 0.03 | 96.26 | 91.03 | 37.68 |
| KD2 | 11394051900 | 11377847400 | 99.86 | 0.03 | 97.34 | 93.02 | 40.18 |
| KE1 | 10455830700 | 10441926000 | 99.87 | 0.03 | 97.73 | 93.47 | 39.48 |
| KE2 | 22805772900 | 22611616800 | 99.15 | 0.04 | 94.14 | 87.2 | 38.63 |
| KF1 | 10703553900 | 10688679000 | 99.86 | 0.03 | 97.78 | 93.56 | 39.55 |
| KF2 | 13262647200 | 13177446000 | 99.36 | 0.03 | 95.97 | 90.64 | 37.66 |
| Kyr | 8945049448 | 8979111710 | 99.38 | 0.03 | 97.14 | 89.88 | 42.57 |
| Rus | 9143354894 | 9174609276 | 99.34 | 0.02 | 98.5 | 92.42 | 38.3 |
| Swe | 13788343410 | 13842519176 | 99.39 | 0.02 | 98.79 | 92.79 | 39.98 |
| Swi1 | 13300134086 | 13364984922 | 99.49 | 0.03 | 97.61 | 90.2 | 40.51 |
| Swi2 | 13926376400 | 13974411124 | 99.34 | 0.02 | 99.31 | 93.93 | 37.55 |
| Taj | 11782210764 | 11824882844 | 99.36 | 0.02 | 99.12 | 93.64 | 36.36 |
| TC1 | 19229094600 | 19163923500 | 99.66 | 0.03 | 96.02 | 90.53 | 38.41 |
| TC2 | 18362369400 | 18298855200 | 99.65 | 0.03 | 96.57 | 91.69 | 38.43 |
| TGG1 | 11160099300 | 11142984300 | 99.85 | 0.03 | 97.71 | 93.7 | 40.04 |
| TGG2 | 11674034700 | 11655866400 | 99.84 | 0.03 | 97.37 | 93.02 | 40.53 |
| TGG3 | 11958380700 | 11940497700 | 99.85 | 0.03 | 97.56 | 93.4 | 39.48 |
| TGG4 | 11868649200 | 11850252300 | 99.84 | 0.03 | 97.27 | 93.01 | 39.9 |
| TGG5 | 11084604600 | 11067224700 | 99.84 | 0.03 | 97.78 | 94.04 | 40.24 |
| Tur1 | 22865317378 | 22920106310 | 99.24 | 0.02 | 98.34 | 92.9 | 36.99 |
| Tur2 | 13949746442 | 13987439470 | 99.27 | 0.02 | 98.76 | 93.33 | 37.53 |
| Tur3 | 8935426978 | 8984988552 | 99.55 | 0.02 | 99.54 | 93.36 | 39.21 |
| UK | 18842955980 | 18903012926 | 99.32 | 0.02 | 99.89 | 94.82 | 36.76 |

**Table S3.**
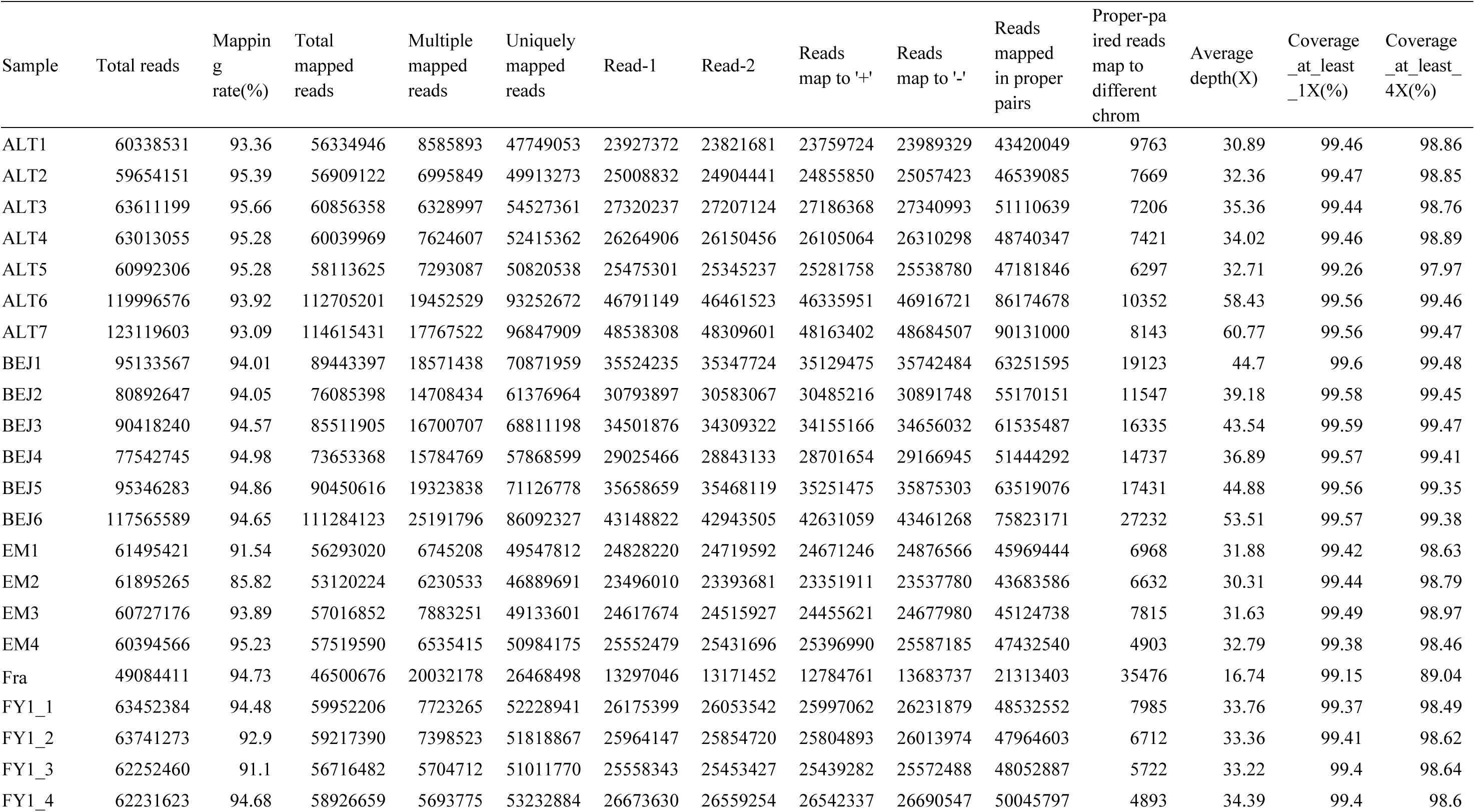
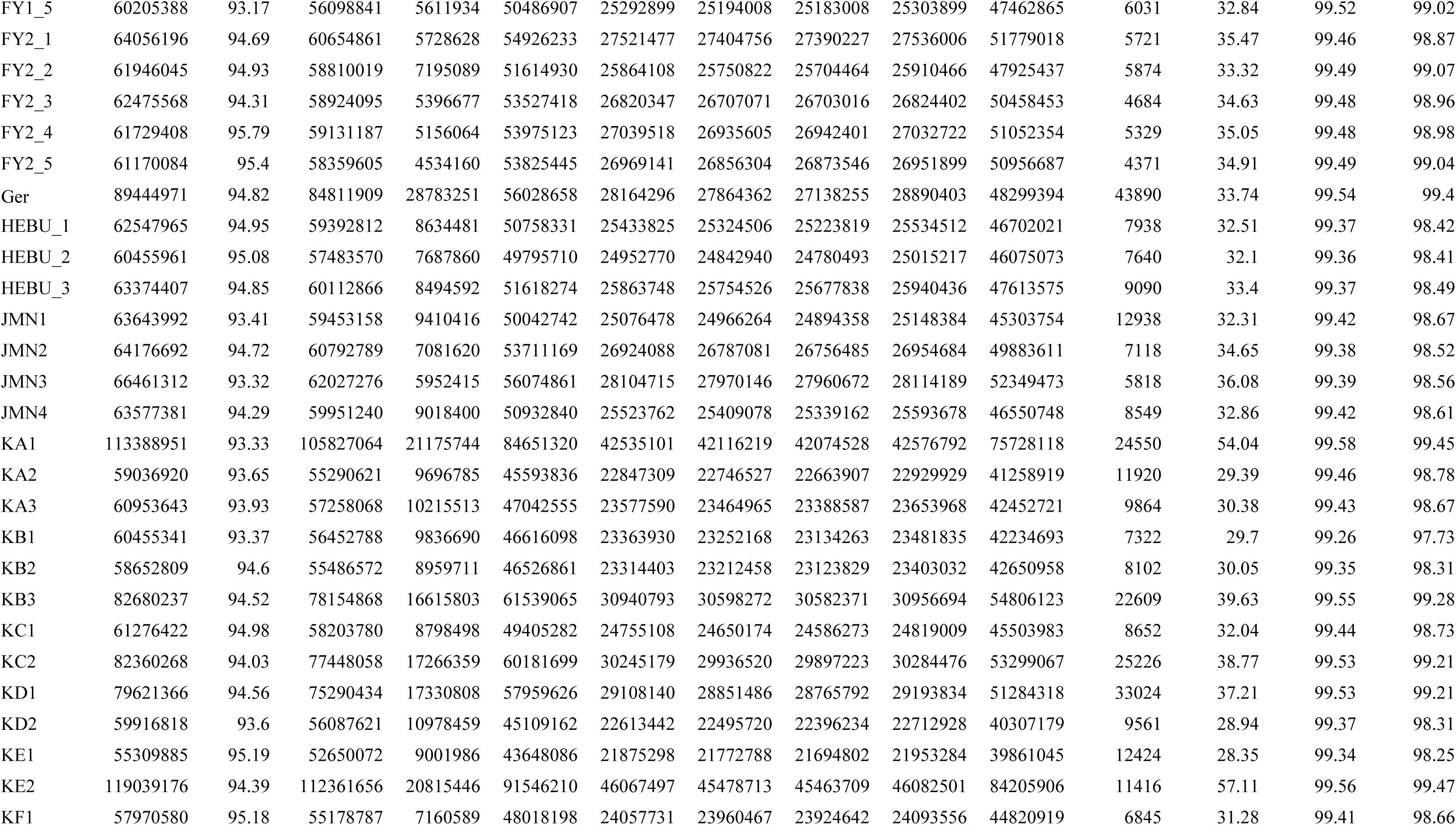
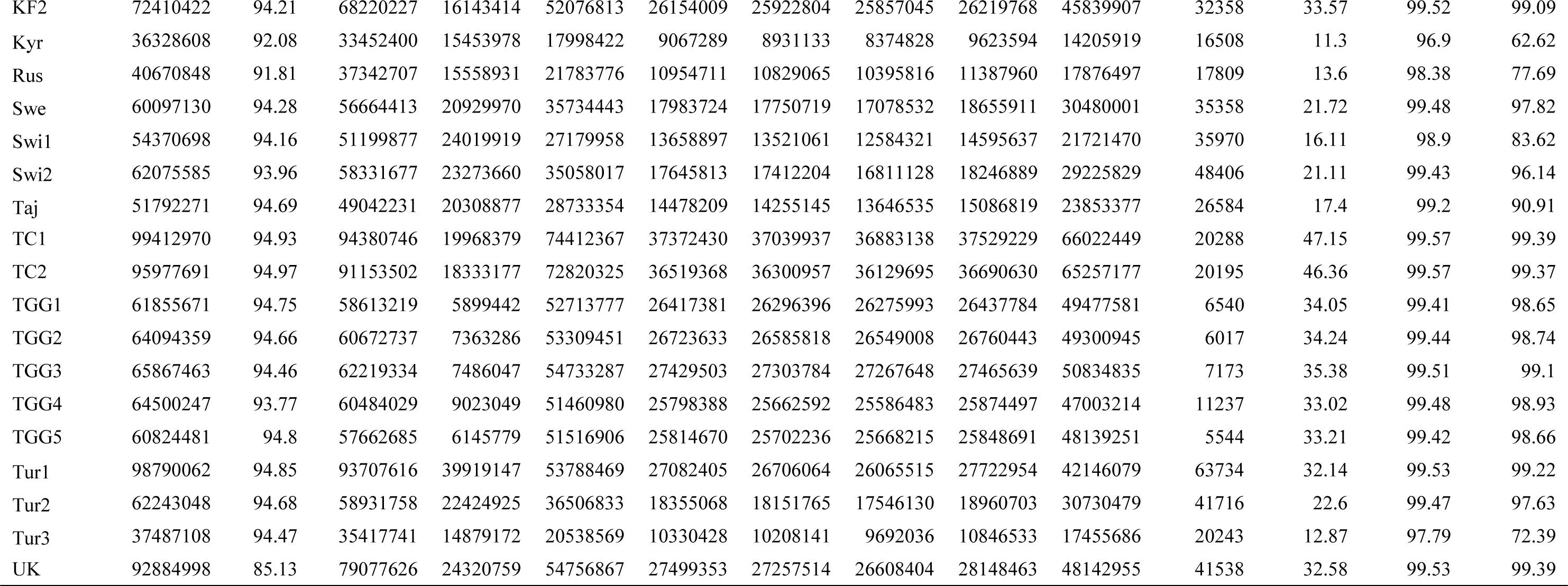
Resequencing Mapping data

**Table S4.**
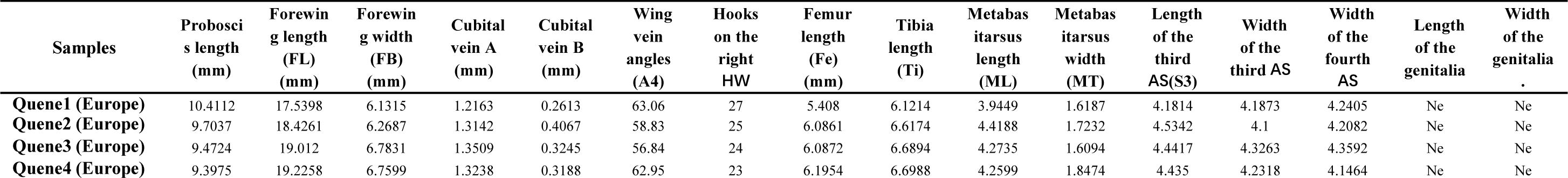
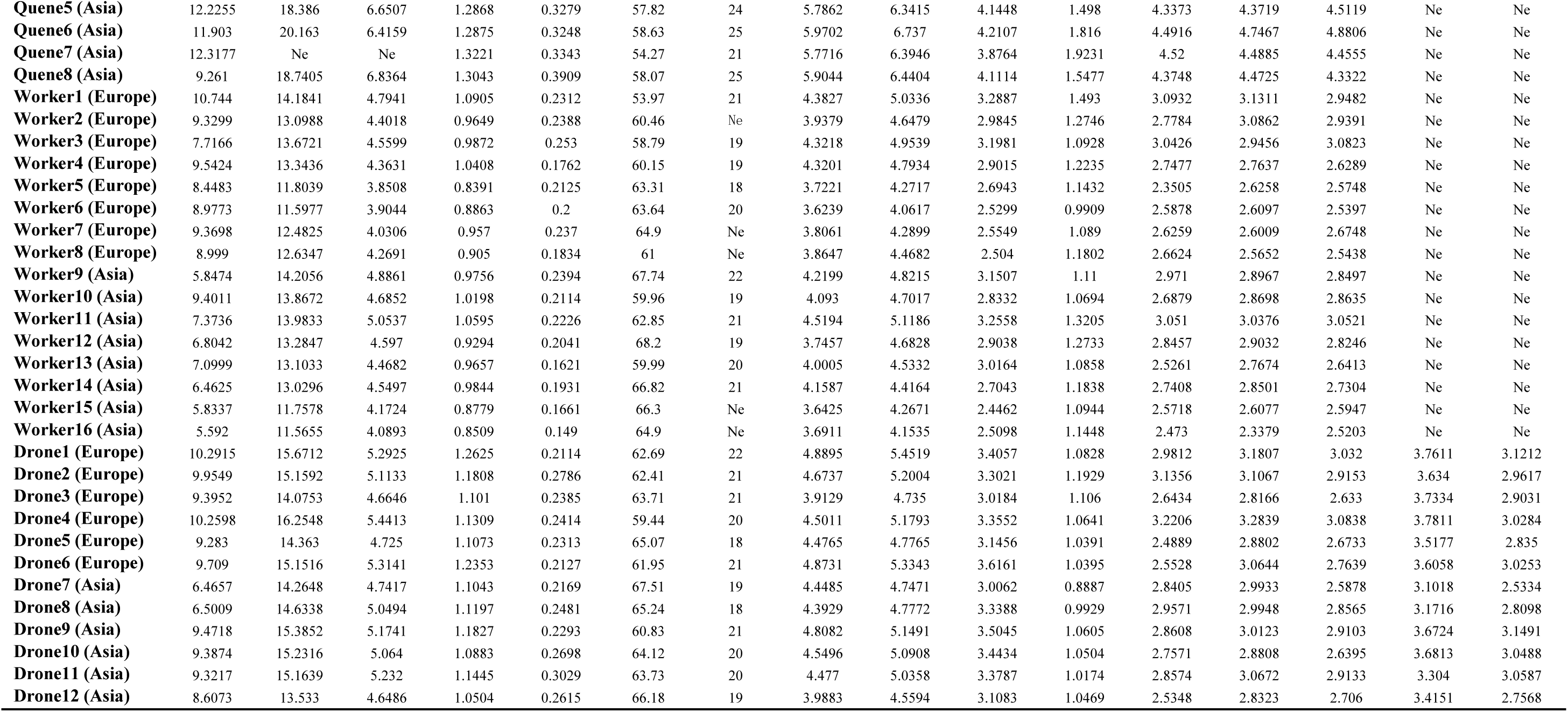
Morphological parameters of *Bombus terrestris* from Asia and Europe

**Table S5.** GO enrichment analysis of Asia samples selected genes

| GO.ID | Term | Annotated | Selected | Expected | Fis p-value |
| --- | --- | --- | --- | --- | --- |
| GO:0007165 | signal transduction | 484 | 34 | 16.95 | 4.10E-05 |
| GO:0007154 | cell communication | 496 | 34 | 17.37 | 6.80E-05 |
| GO:0023052 | signaling | 498 | 34 | 17.44 | 7.40E-05 |
| GO:0050789 | regulation of biological process | 994 | 55 | 34.81 | 0.00012 |
| GO:0050794 | regulation of cellular process | 964 | 53 | 33.76 | 0.0002 |
| GO:0050896 | response to stimulus | 644 | 39 | 22.56 | 0.00028 |
| GO:0065007 | biological regulation | 1042 | 55 | 36.49 | 0.00043 |
| GO:0051716 | cellular response to stimulus | 598 | 36 | 20.94 | 0.00057 |
| GO:0043087 | regulation of GTPase activity | 17 | 4 | 0.6 | 0.00241 |
| GO:0035556 | intracellular signal transduction | 193 | 15 | 6.76 | 0.00281 |
| GO:0006487 | protein N-linked glycosylation | 4 | 2 | 0.14 | 0.00698 |
| GO:0016477 | cell migration | 4 | 2 | 0.14 | 0.00698 |
| GO:0051336 | regulation of hydrolase activity | 23 | 4 | 0.81 | 0.00762 |
| GO:0007166 | cell surface receptor signaling pathway | 69 | 7 | 2.42 | 0.00985 |
| GO:0007275 | multicellular organism development | 39 | 5 | 1.37 | 0.01087 |
| GO:0006182 | cGMP biosynthetic process | 6 | 2 | 0.21 | 0.01666 |
| GO:0007093 | mitotic cell cycle checkpoint | 6 | 2 | 0.21 | 0.01666 |
| GO:0045930 | negative regulation of mitotic cell cycl... | 6 | 2 | 0.21 | 0.01666 |
| GO:0046068 | cGMP metabolic process | 6 | 2 | 0.21 | 0.01666 |
| GO:0048870 | cell motility | 6 | 2 | 0.21 | 0.01666 |
| GO:0051674 | localization of cell | 6 | 2 | 0.21 | 0.01666 |
| GO:0016051 | carbohydrate biosynthetic process | 16 | 3 | 0.56 | 0.01687 |
| GO:0048856 | anatomical structure development | 46 | 5 | 1.61 | 0.02128 |
| GO:0040011 | locomotion | 7 | 2 | 0.25 | 0.0228 |
| GO:0051304 | chromosome separation | 8 | 2 | 0.28 | 0.0297 |
| GO:0032502 | developmental process | 52 | 5 | 1.82 | 0.03417 |
| GO:0043547 | positive regulation of GTPase activity | 9 | 2 | 0.32 | 0.03732 |
| GO:0052652 | cyclic purine nucleotide metabolic proce... | 9 | 2 | 0.32 | 0.03732 |
| GO:0007155 | cell adhesion | 38 | 4 | 1.33 | 0.04241 |
| GO:0022610 | biological adhesion | 38 | 4 | 1.33 | 0.04241 |
| GO:0009653 | anatomical structure morphogenesis | 10 | 2 | 0.35 | 0.0456 |
| GO:0051345 | positive regulation of hydrolase activit... | 10 | 2 | 0.35 | 0.0456 |
| GO:0048523 | negative regulation of cellular process | 57 | 5 | 2 | 0.04797 |

**Table S6.** KEGG enrichment analysis of Asia samples selected genes

| KEGG Term | ID | Input number | Background number | P-Value |
| --- | --- | --- | --- | --- |
| MAPK signaling pathway - fly | bter04013 | 10 | 82 | 3.73E-05 |
| Wnt signaling pathway | bter04310 | 9 | 69 | 5.69E-05 |
| Glycerophospholipid metabolism | bter00564 | 5 | 57 | 0.012256 |
| Apoptosis - fly | bter04214 | 4 | 46 | 0.025026 |
| Hippo signaling pathway - fly | bter04391 | 4 | 48 | 0.028428 |

**Table S7.**
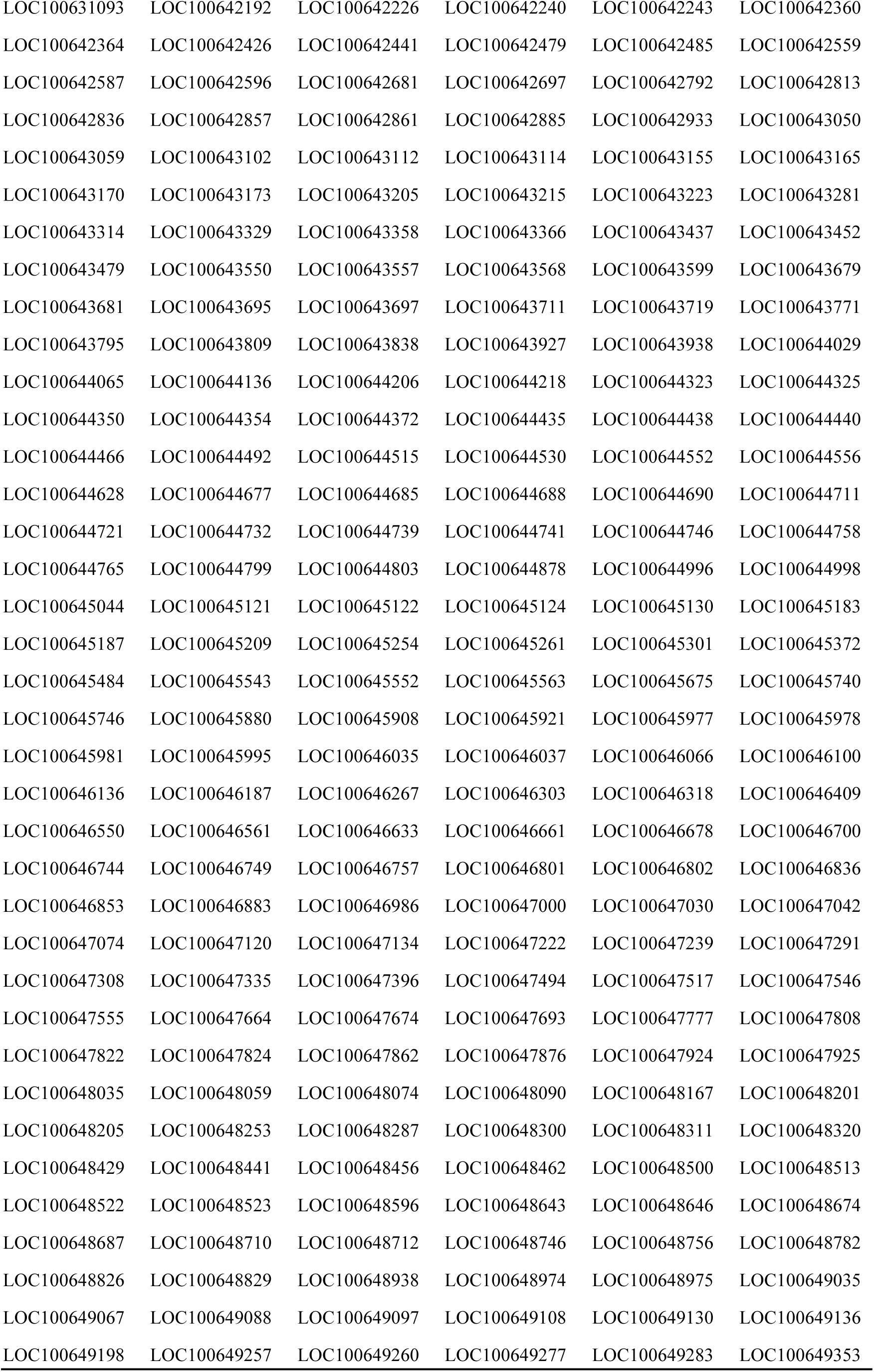
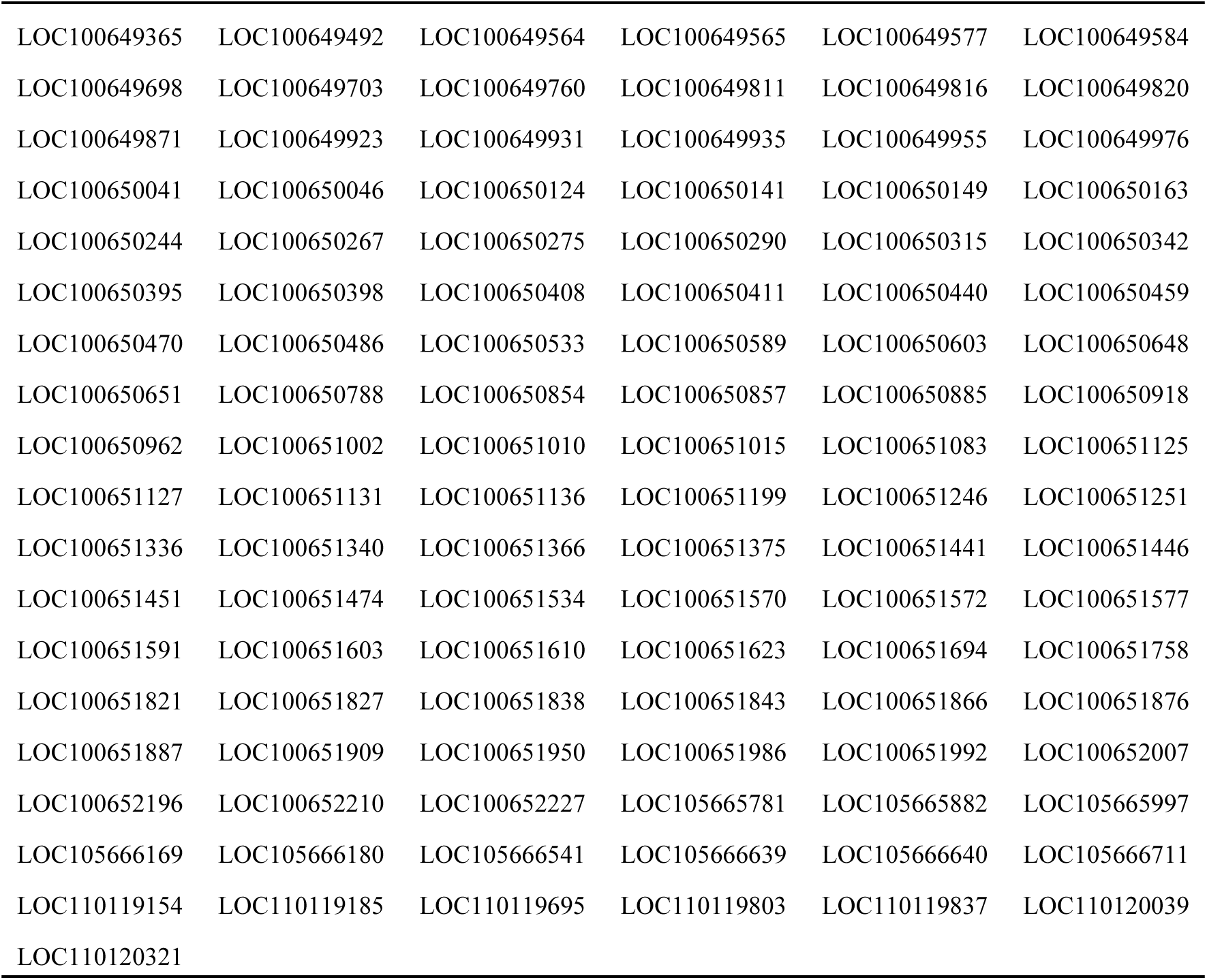
Asia samples selected genes

|  |  |  |  |  |  |
| --- | --- | --- | --- | --- | --- |
| LOC100631093 | LOC100642192 | LOC100642226 | LOC100642240 | LOC100642243 | LOC100642360 |
| LOC100642364 | LOC100642426 | LOC100642441 | LOC100642479 | LOC100642485 | LOC100642559 |
| LOC100642587 | LOC100642596 | LOC100642681 | LOC100642697 | LOC100642792 | LOC100642813 |
| LOC100642836 | LOC100642857 | LOC100642861 | LOC100642885 | LOC100642933 | LOC100643050 |
| LOC100643059 | LOC100643102 | LOC100643112 | LOC100643114 | LOC100643155 | LOC100643165 |
| LOC100643170 | LOC100643173 | LOC100643205 | LOC100643215 | LOC100643223 | LOC100643281 |
| LOC100643314 | LOC100643329 | LOC100643358 | LOC100643366 | LOC100643437 | LOC100643452 |
| LOC100643479 | LOC100643550 | LOC100643557 | LOC100643568 | LOC100643599 | LOC100643679 |
| LOC100643681 | LOC100643695 | LOC100643697 | LOC100643711 | LOC100643719 | LOC100643771 |
| LOC100643795 | LOC100643809 | LOC100643838 | LOC100643927 | LOC100643938 | LOC100644029 |
| LOC100644065 | LOC100644136 | LOC100644206 | LOC100644218 | LOC100644323 | LOC100644325 |
| LOC100644350 | LOC100644354 | LOC100644372 | LOC100644435 | LOC100644438 | LOC100644440 |
| LOC100644466 | LOC100644492 | LOC100644515 | LOC100644530 | LOC100644552 | LOC100644556 |
| LOC100644628 | LOC100644677 | LOC100644685 | LOC100644688 | LOC100644690 | LOC100644711 |
| LOC100644721 | LOC100644732 | LOC100644739 | LOC100644741 | LOC100644746 | LOC100644758 |
| LOC100644765 | LOC100644799 | LOC100644803 | LOC100644878 | LOC100644996 | LOC100644998 |
| LOC100645044 | LOC100645121 | LOC100645122 | LOC100645124 | LOC100645130 | LOC100645183 |
| LOC100645187 | LOC100645209 | LOC100645254 | LOC100645261 | LOC100645301 | LOC100645372 |
| LOC100645484 | LOC100645543 | LOC100645552 | LOC100645563 | LOC100645675 | LOC100645740 |
| LOC100645746 | LOC100645880 | LOC100645908 | LOC100645921 | LOC100645977 | LOC100645978 |
| LOC100645981 | LOC100645995 | LOC100646035 | LOC100646037 | LOC100646066 | LOC100646100 |
| LOC100646136 | LOC100646187 | LOC100646267 | LOC100646303 | LOC100646318 | LOC100646409 |
| LOC100646550 | LOC100646561 | LOC100646633 | LOC100646661 | LOC100646678 | LOC100646700 |
| LOC100646744 | LOC100646749 | LOC100646757 | LOC100646801 | LOC100646802 | LOC100646836 |
| LOC100646853 | LOC100646883 | LOC100646986 | LOC100647000 | LOC100647030 | LOC100647042 |
| LOC100647074 | LOC100647120 | LOC100647134 | LOC100647222 | LOC100647239 | LOC100647291 |
| LOC100647308 | LOC100647335 | LOC100647396 | LOC100647494 | LOC100647517 | LOC100647546 |
| LOC100647555 | LOC100647664 | LOC100647674 | LOC100647693 | LOC100647777 | LOC100647808 |
| LOC100647822 | LOC100647824 | LOC100647862 | LOC100647876 | LOC100647924 | LOC100647925 |
| LOC100648035 | LOC100648059 | LOC100648074 | LOC100648090 | LOC100648167 | LOC100648201 |
| LOC100648205 | LOC100648253 | LOC100648287 | LOC100648300 | LOC100648311 | LOC100648320 |
| LOC100648429 | LOC100648441 | LOC100648456 | LOC100648462 | LOC100648500 | LOC100648513 |
| LOC100648522 | LOC100648523 | LOC100648596 | LOC100648643 | LOC100648646 | LOC100648674 |
| LOC100648687 | LOC100648710 | LOC100648712 | LOC100648746 | LOC100648756 | LOC100648782 |
| LOC100648826 | LOC100648829 | LOC100648938 | LOC100648974 | LOC100648975 | LOC100649035 |
| LOC100649067 | LOC100649088 | LOC100649097 | LOC100649108 | LOC100649130 | LOC100649136 |
| LOC100649198 | LOC100649257 | LOC100649260 | LOC100649277 | LOC100649283 | LOC100649353 |
| LOC100649365 | LOC100649492 | LOC100649564 | LOC100649565 | LOC100649577 | LOC100649584 |
| LOC100649698 | LOC100649703 | LOC100649760 | LOC100649811 | LOC100649816 | LOC100649820 |
| LOC100649871 | LOC100649923 | LOC100649931 | LOC100649935 | LOC100649955 | LOC100649976 |
| LOC100650041 | LOC100650046 | LOC100650124 | LOC100650141 | LOC100650149 | LOC100650163 |
| LOC100650244 | LOC100650267 | LOC100650275 | LOC100650290 | LOC100650315 | LOC100650342 |
| LOC100650395 | LOC100650398 | LOC100650408 | LOC100650411 | LOC100650440 | LOC100650459 |
| LOC100650470 | LOC100650486 | LOC100650533 | LOC100650589 | LOC100650603 | LOC100650648 |
| LOC100650651 | LOC100650788 | LOC100650854 | LOC100650857 | LOC100650885 | LOC100650918 |
| LOC100650962 | LOC100651002 | LOC100651010 | LOC100651015 | LOC100651083 | LOC100651125 |
| LOC100651127 | LOC100651131 | LOC100651136 | LOC100651199 | LOC100651246 | LOC100651251 |
| LOC100651336 | LOC100651340 | LOC100651366 | LOC100651375 | LOC100651441 | LOC100651446 |
| LOC100651451 | LOC100651474 | LOC100651534 | LOC100651570 | LOC100651572 | LOC100651577 |
| LOC100651591 | LOC100651603 | LOC100651610 | LOC100651623 | LOC100651694 | LOC100651758 |
| LOC100651821 | LOC100651827 | LOC100651838 | LOC100651843 | LOC100651866 | LOC100651876 |
| LOC100651887 | LOC100651909 | LOC100651950 | LOC100651986 | LOC100651992 | LOC100652007 |
| LOC100652196 | LOC100652210 | LOC100652227 | LOC105665781 | LOC105665882 | LOC105665997 |
| LOC105666169 | LOC105666180 | LOC105666541 | LOC105666639 | LOC105666640 | LOC105666711 |
| LOC110119154 | LOC110119185 | LOC110119695 | LOC110119803 | LOC110119837 | LOC110120039 |
| LOC110120321 |  |  |  |  |  |

